# *Bona fide* cytotoxic iNKT cells with superior antitumour responses identified in mice and humans

**DOI:** 10.1101/2024.07.18.603987

**Authors:** Carolina de Amat Herbozo, Stephanie WY Wong, Meggie Kuypers, Guy Ilango, Zhewei Liu, Jessica A Mathews, Albert Nguyen, André Veillette, Jennifer L Gommerman, Thomas Baranek, Sarah Q Crome, Ming Sound Tsao, Adrian G Sacher, Christophe Paget, Thierry Mallevaey

## Abstract

Adoptive cell therapies (ACT) using unmodified or engineered invariant Natural Killer T (iNKT) cells are in clinical trials for cancer treatment. While promising, outcomes are still suboptimal, possibly due to iNKT cell heterogeneity. This study identified unique iNKT cell populations with strong cytotoxic activity in mice and humans. In mice, iNKT1c cells showed potent CD1d-dependent and -independent antitumor functions, with responses influenced by NK receptors. IL-15 enhances their cytolytic activity, while retinoic acid is essential for their generation. In humans, terminally differentiated CD57^+^ iNKT cells showed marked NK-receptor expression and most effectively killed C1R tumor cells *in vitro*. These cells appeared activated/exhausted within tumors of non-small cell lung cancer patients. Additionally, a central memory-like CD62L^+^ CD4^-^ iNKT subset efficiently expanded and generated cytotoxic CD57^+^ iNKT cells *in vitro*, making them an appealing candidate for ACT. This study paves the way for the design of more effective iNKT cell-based immunotherapies.

## INTRODUCTION

Invariant Natural Killer T (iNKT) cells are unconventional T cells that respond to glycolipid antigens, such as α-galactosylceramide (αGalCer), presented by the major histocompatibility complex (MHC) class Ib molecule CD1d ^1,2^. Given their rapid response and functional heterogeneity, iNKT cells play crucial protective functions in a variety of diseases, including cancer ^3–8^. Pre-clinical studies have demonstrated their antitumor therapeutic potential against hematological malignancies and solid tumors ^9–12^. Moreover, since CD1d is non-polymorphic, iNKT cells are excellent candidates for allogeneic off-the-shelf adoptive cell therapy (ACT). Clinical trials have established that the autologous and allogeneic adoptive transfer of iNKT cells is safe and well tolerated in cancer patients ^13–19^. However, the response rate and efficacy of iNKT cell-based immunotherapies remain suboptimal, possibly due to the functional heterogeneity of iNKT cells. Therefore, a better understanding of iNKT cell effector subsets is required to harness their full therapeutic potential.

Cytotoxicity is an essential component of antitumor immune responses as it allows for the rapid destruction of tumor cells. Cytotoxic CD8^+^ T cells and NK cells are the *bona fide* cytotoxic populations of the conventional αβ T cells and innate lymphoid lineages, respectively. Due to their potent tumor killing abilities, both populations are highly relevant for ACT ^20–22^. Although iNKT cells can also kill tumor cells ^23–26^, their cytotoxicity is less well characterized than their helper functions. Specifically, it remains unclear whether cytotoxicity is a universal function of all iNKT cells or if it is restricted to one or more dedicated cytotoxic iNKT cell subset(s).

In mice, the three main iNKT cell effector subsets, iNKT1, iNKT2 and iNKT17, are currently defined by their helper functions, which parallel that of T helped (Th) 1, Th2 and Th17 CD4^+^ T cells, respectively ^27,28^. More recently, we and others reported heterogeneity within thymic iNKT1 cells using single cell RNA sequencing (scRNA-seq) ^29^ and we identified three distinct iNKT1 cell clusters: iNKT1a, iNKTb, and iNKT1c. Gene profile and pseudotime analysis suggested that iNKT1a cells may be developmental intermediates. iNKT1b cells expressed high levels of *Nkg7*, *Il2rb*, *Cxcr3*, *Id2* transcripts and transcripts related to Th1 differentiation such as members of the GTPases of the immunity-associated protein (GIMAP) family (*Gimap1*, *3*, *4*, *5* and *6*). Finally, iNKT1c cells were characterized by a cytotoxic gene profile enriched for genes encoding for killer cell lectin-like receptors (including *Klra9* and *Klrk1*), *Fcer1g* (subunit γ of the high affinity IgE receptor), *Gzma*, and *Gzmb* ^29^. These findings were corroborated by another study ^30^. In humans, recent scRNA-seq analyses identified a cluster of peripheral blood iNKT cells characterized by higher expression of genes related to cytotoxicity such as *KLRD1*, *GZMB*, and *GNLY* ^31,32^. However, their cytotoxic activity towards tumor cells and whether they participate in antitumor immune responses has not been determined.

Identifying cytotoxic iNKT cell populations with antitumor activity and understanding the factors that regulate their response is a crucial step to potentiate iNKT cell-based ACTs. In this study, we identified mouse iNKT1b and iNKT1c cell populations in peripheral tissues as well as within melanoma tumors. We further define their transcriptomic profile and demonstrate the cytotoxic activity and potent antitumor responses of iNKT1c cells. iNKT1c cells killed B16F10 melanoma cells in CD1d-dependent and -independent manners. Furthermore, we found that cytotoxic iNKT1c cell responses are regulated by antigen-independent signals such as cytokines and NK receptors. We further describe a role for retinoic acid (RA) signaling in supporting generation or maintenance of iNKT1c cells. In humans, we identified CD57-expressing iNKT cells with strong cytotoxic profiles and tumor killing activity. Importantly, cytotoxic CD57^+^ iNKT cells infiltrated solid tumors in non-small cell lung cancer (NSCLC) patients, where they appeared to display hallmarks of activation and exhaustion. Moreover, through pseudotime analysis and *in vitro* cell expansion of peripheral iNKT cells we identified a central memory-like iNKT cell population capable of generating cytotoxic iNKT cells, making it a promising candidate for ACT. Altogether, our findings define highly cytotoxic iNKT populations in mice and humans and demonstrate the potential of this iNKT cell subset for cancer immunotherapy.

## RESULTS

### Two distinct iNKT1 populations are found in mouse tissues

Our previous single cell RNA sequencing (scRNA-seq) analysis of mouse thymic iNKT cells revealed two main clusters of iNKT1 cells, namely iNKT1b and iNKT1c, with transcriptomic profiles reminiscent of helper and cytotoxic functions, respectively^29^. We used the markers defined in this previous study to detect iNKT1b and iNKT1c cells by fluorescence-assisted cell sorting (FACS) in the thymus and peripheral tissues. Specifically, after excluding CD24^+^ (iNKT0) and RORγt^+^ or Sdc-1^+^ iNKT17 cells, iNKT1b cells were defined as NK1.1^+^ Sca-1^+^ cells, while iNKT1c cells were gated as NK1.1^+^ Sca-1^-^ (**Fig. 1A**). To corroborate that the gated populations correspond to the previously identified transcriptomic clusters, we analyzed the gene expression profiles of FACS-sorted T-bet^+^ NK1.1^+^ Sca-1^+^ iNKT1b and T-bet^+^ NK1.1^+^ Sca-1^-^ iNKT1c from the thymus, spleen, and liver of T-bet-RFP reporter mice, using bulk RNA sequencing (RNA-seq) (**Sup. Fig. 1A**). Principal component analysis (PCA) revealed that iNKT1b cells segregated from iNKT1c cells in all tissues along PC1 and PC2, which accounted for a total of 83% of the variation. The segregation between iNKT1b and iNKT1c cells was more pronounced in the spleen and liver, and they clustered apart from the thymic populations, suggesting that tissue-specific signals may also impact gene expression (**Fig. 1B**). We performed gene set enrichment analysis (GSEA) ^33,34^ with custom gene sets curated from our previous thymic scRNA-seq data to establish iNKT1b (upregulated in iNKT1b compared to iNKT1c) and iNKT1c (downregulated in iNKT1b respective to iNKT1c) gene signatures. In all tissues, a significant enrichment score was observed for their respective population, indicating that the iNKT1b and iNKT1c populations identified by FACS and scRNA-seq have matching gene signatures (**Fig. 1C**). Specifically, and in agreement with the single cell transcriptomic data, iNKT1b cells showed higher expression of the genes *Gimap3/4/6*, *Plac8*, *Icos*, *Ramp1*, *Cd4*, and *Il4*, while iNKT1c cells were characterized by the increased expression of *Fcer1g*, *Ccl5*, *Gzma/b*, *Cd244a*, *Klf2*, *S100a4*, *Vim*, killer cell lectin-like receptor genes (e.g., *Klra1/5/6/7/9*, *Klre1*), *Crip1*, and *Lgals1* (**Fig. 1D**, **Sup. Fig. 1B**). As expected, *Ly6a* (encoding for Sca-1) was highly expressed by iNKT1b cells. Some of the differentially expressed genes (DEGs) were predominant in the thymus (e.g., *Cd244a*, *Klra5*, *Ccr10*), while others were enriched in peripheral tissues (e.g., *Ncr1*, *Ccl5*, *Gzmm*) (**Fig. 1D**). We confirmed the higher expression of GzmA, GzmB, 2B4 (CD244), and Klf2 by iNKT1c cells using FACS (**Fig. 1E, F**). The iNKT1c signature was also consistent with the thymic and lung C2 cytotoxic iNKT cell population recently described by the Ikuta group ^35^. In addition to iNKT1b and iNKT1c subsets, peripheral tissues contained a sizeable population of T-bet^+^ NK1.1^-^ iNKT1 cells (**Sup. Fig. 1B**), which we included in our transcriptomic analysis. The Euclidean distance calculation and PCA revealed that peripheral T-bet^+^ NK1.1^-^ iNKT1 cells and iNKT1b cells are highly similar (**Sup. Fig. 1C, D**). DEGs significantly upregulated in T-bet^+^ NK1.1^-^ iNKT1 cells compared to iNKT1b cells were related to cell division, including cyclins (e.g., *Ccnb1*, *Ccnf*), cyclin-dependent kinases (e.g., *Cdk1*, *Cdkn3*), cell division cycle associated genes (e.g., *Cdca3*, *Cdc6*), kinesin family members (e.g., *Kif11*, *Kif2c*), kinetochore complex components (e.g., *Spc24*, *Spc25*), and histones (e.g., *H2ac8*, *H2bc11*) (**Sup. Fig. 1E**). This was confirmed through gene ontology (GO) enrichment analysis (**Sup. Fig. 1F**). On the other hand, *Mmp9*, *Hhex*, and some killer cell lectin-like receptor genes were downregulated in T-bet^+^ NK1.1^-^ iNKT1 cells compared to iNKT1b cells (**Sup. Fig. 1E**). This suggests that the T-bet^+^ NK1.1^-^ iNKT1 cells may correspond to a fraction of iNKT1b cells undergoing active division. Therefore, we focused on NK1.1^+^ iNKT1b and iNKT1c cells for the rest of this study. Both populations were present in all peripheral tissues tested, whereby iNKT1b cells dominated and iNKT1c cells represented no more than 10% of total iNKT cells (**Fig. 1G**). However, when we established B16F10 tumors in the flank of C57BL/6 (B6) mice, we found similar proportions of iNKT1b and iNKT1c within tumor-infiltrating iNKT cells (**Fig. 1H**). Together, these data demonstrate that iNKT1b (helper profile) and iNKT1c (cytotoxic profile) cells are present in both the thymus as well as peripheral tissues, with a preferential accumulation of iNKT1c cells in B16F10 melanoma tumors.

**Figure 1.**
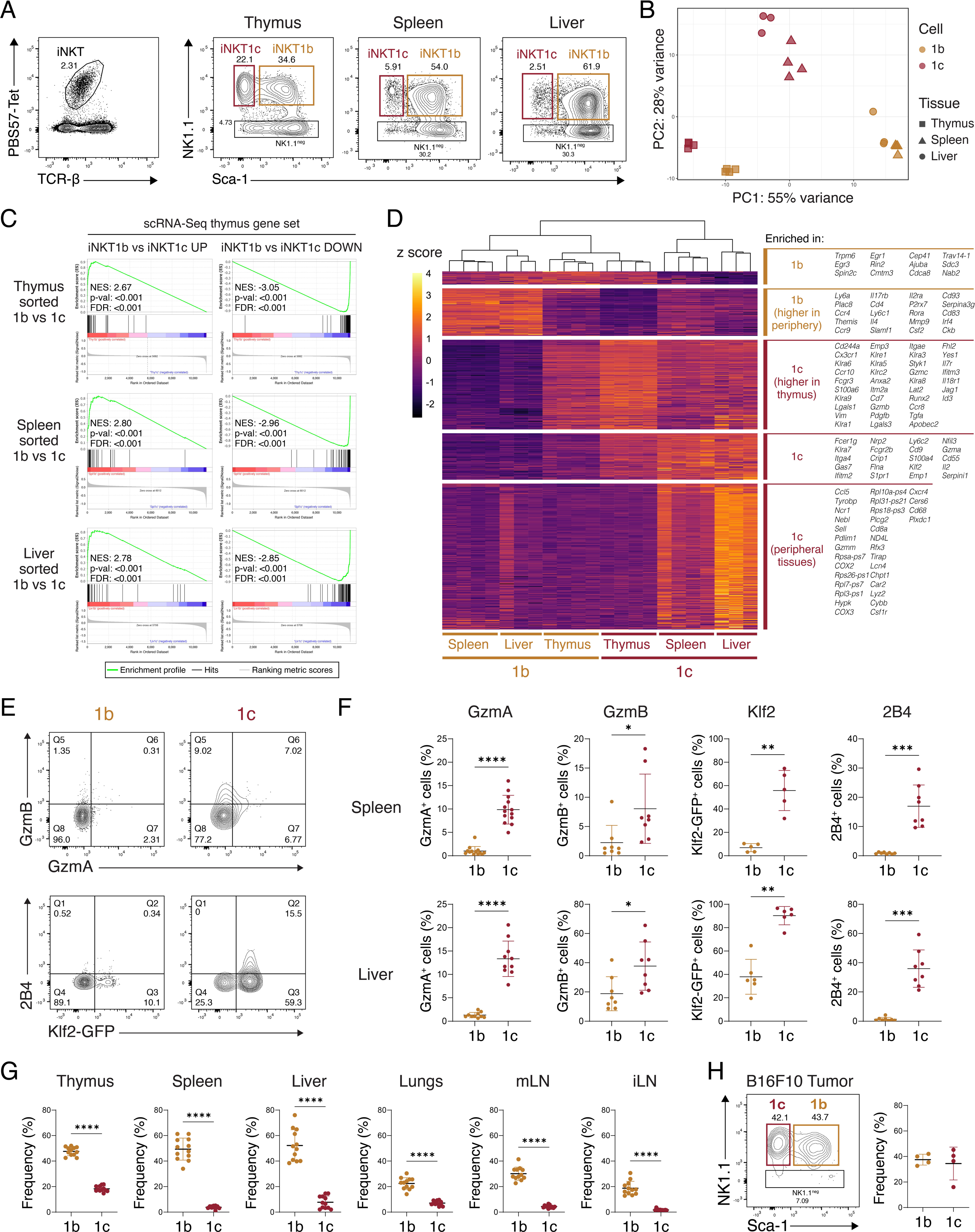
Two subsets of iNKT1 cells are found in the thymus and peripheral tissues. **A)** Gating strategy for iNKT1 populations in the thymus, spleen, and liver from B6 mice. After gating on iNKT cells, CD24^+^ and RORγt^+^ cells are excluded before gating on NK1.1^+^ Sca-1^+^ iNKT1b (1b) cells and NK1.1^+^ Sca-1^-^ iNKT1c (1c) cells. **B)** PCA plot of sorted iNKT1b and iNKT1c cells analyzed by bulk RNA-seq. **C)** GSEA plots showing the enrichment scores for gene sets curated from our published scRNA-seq data of thymic iNKT cells. **D)** Heatmap showing DEGs (log2 fold change ≥ 1, adjusted p-value < 0.05) from thymic and peripheral iNKT1b and iNKT1c cells. **E, F)** Representative FACS plots (E) and the frequency (F) of iNKT1b and iNKT1c cells expressing GzmA, GzmB, Klf2 and 2B4. **G)** Frequency of iNKT1b and iNKT1c populations from total iNKT cells in the indicated tissues. **H)** Representative FACS plot (left) and frequencies (right) of iNKT1b and iNKT1c cells infiltrating subcutaneous B16F10 tumors established in B6 mice. Data shown the mean values ± SD. **** p<0.0001, *** p<0.001, ** p<0.01, * p<0.05, Mann-Whitney test.

### Higher granzyme response and specific killing activity by iNKT1c cells

Although the proportion of GzmA/B^+^ cells was higher in iNKT1c cells than in iNKT1b cells, overall granzyme expression was low at steady state. However, GzmA was highly expressed in splenic and hepatic iNKT1c cells 3, 7, and 10 days following the intravenous injection of αGalCer-loaded bone marrow-derived dendritic cells (BMDC) *in vivo*, confirming that iNKTc cells display a cytotoxic program following activation (**Fig. 2A, B**). GzmA was induced to a lower extent in iNKT1b cells and returned to basal levels 10 days post-injection of αGalCer-loaded BMDCs (**Fig. 2A, B**). As reported in previous studies, antigenic stimulation did not enhance GzmB expression by iNKT cells at any time point following αGalCer-loaded BMDC injection (**Fig. 2A** and not shown) ^36^. On the other hand, *in vitro* stimulation of splenocytes with PMA and ionomycin or IL-15 increased the proportion of GzmB^+^ iNKT1c cells, and to a lower extent iNKT1b cells (**Fig. 2C**). GzmB expression by IL-15-stimulated iNKT1b and iNKT1c cells was not impaired by anti-CD1d neutralizing antibodies (**Fig. 2D**). As a control, we showed that anti-CD1d neutralizing antibodies severely blunted TCR signaling upon αGalCer-mediated stimulation, as assessed by Nur77 expression (**Fig. 2D**). This suggests that IL-15 induces GzmB expression by iNKT1b and iNKT1c cells independently of TCR engagement.

**Figure 2.**
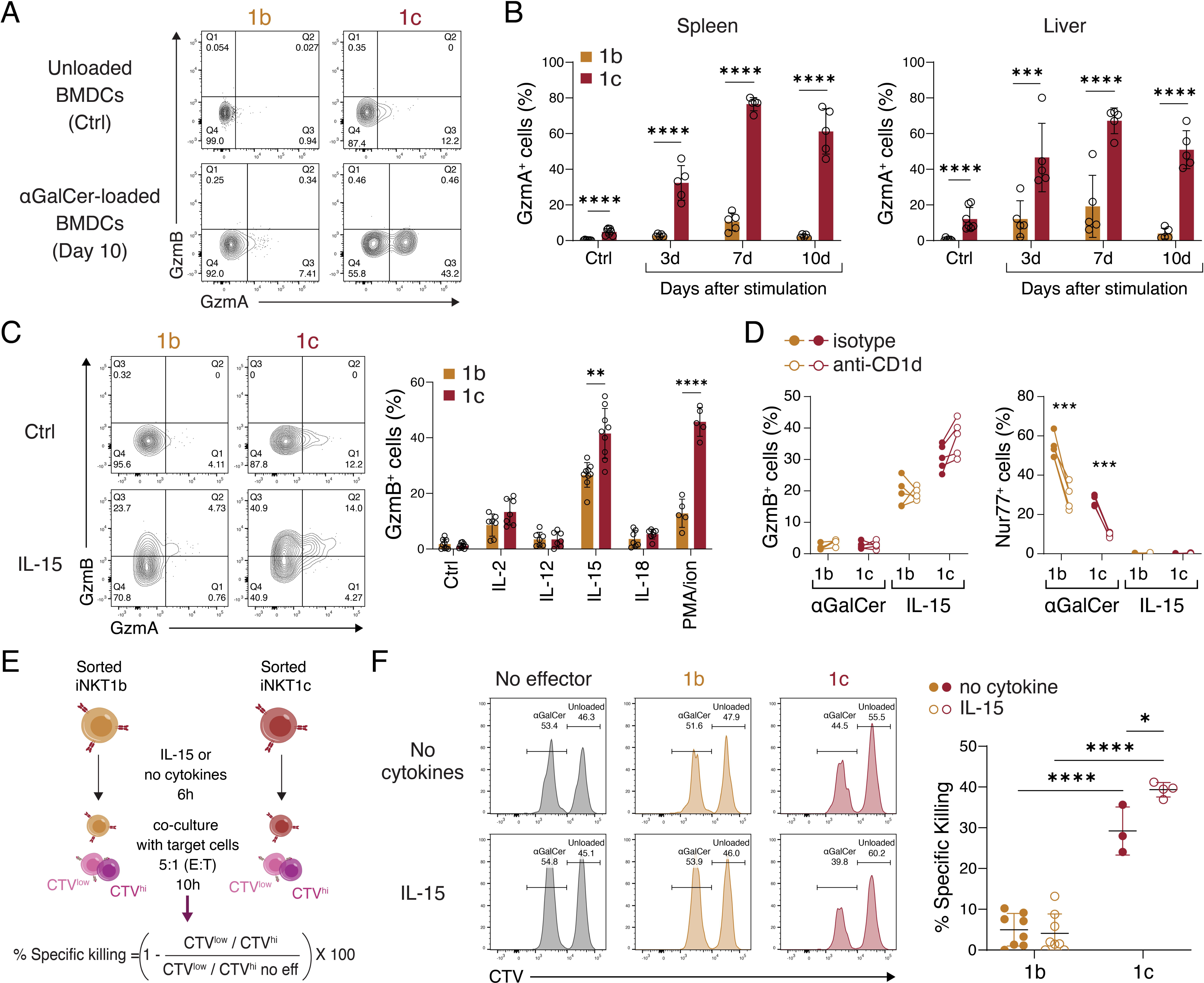
Cytotoxicity induction and superior killing activity of iNKT1c cells. **A, B)** B6 mice were injected intravenously with αGalCer-loaded or unloaded (Ctrl) BMDCs. The phenotype of iNKT cells from the spleen and liver was evaluated by FACS at different time points. Representative plots (A) and frequencies (B) of GzmA^+^ cells within iNKT1b (1b) and iNKT1c (1c) cells. **C)** B6 splenocytes were stimulated *in vitro* with the indicated cytokines (100 ng/mL), PMA and ionomycin (PMA/ion), or left unstimulated (Ctrl). Representative plots showing the expression of GzmA/B (left) and the frequency of GzmB^+^ iNKT1b and iNKT1c cells (right) are shown. **D)** Splenocytes were stimulated with IL-15 (100 ng/mL) or αGalCer (75 ng/mL) in presence of anti-CD1d (clone 1B1) or isotype control. The frequency of GzmB-expressing iNKT1b and iNKT1c cells is shown for each condition. **E)** αGalCer-loaded CD1d-GFP-transduced EL4 cells and unloaded GFP-transduced EL4 cells were stained with CellTrace Violet at 0.25 μM (CTV^low^) and 2.5 μM (CTV^hi^), respectively. Then a 1:1 mix of these target cells were co-cultured with iNKT1b and iNKT1c cells purified from the liver of B6 mice. Effector iNKT1b and iNKT1c cells were pre-incubated with IL-15 (100 ng/mL) or left unstimulated for 6 h before the co-culture. Schematic of the experiment and formula to calculate the percentage of specific killing is shown. **F)** Representative histograms showing the peaks of the respective target cells after the co-culture (left) and percentage of specific killing in each condition (right) (CV<15%). Data represents mean values ± SD. **** p<0.0001, *** p<0.001, ** p<0.01, * p<0.05, 2-way ANOVA of rank-transformed data with Tukey correction for multiple comparison (B, F) and multiple t-test with Holm-Šídák correction (C, D).

Next, we directly assessed the cytotoxicity of iNKT1b and iNKT1c populations *in vitro* against αGalCer-loaded CD1d-GFP EL4 lymphoma target cells (transduced to overexpress CD1d) relative to control unloaded EL4-GFP cells, labeled with two different concentrations of cell trace violet (CTV) (**Fig. 2E**). iNKT1c cells killed αGalCer-loaded CD1d-expressing EL4 cells with greater efficiency than iNKT1b cells (**Fig. 2F**). In addition, consistent with its effect on GzmB production, IL-15 stimulation increased the specific killing activity of iNKT1c cells, but not of iNKT1b cells. (**Fig. 2F**). These results demonstrate that iNKT1c cells exert higher cytotoxic activity than iNKT1b, which is potentiated by IL-15.

### iNKT1c cells have weaker cytokine responses to the cognate antigen αGalCer

Our RNA-seq analyses revealed that iNKT1b and iNKT1c cells differed in their expression of cytokine genes, with *Ifng*, *Il2*, and *Flt3l* transcripts enriched in peripheral iNKT1c cells, and transcripts such as *Il4*, *Csf2*, *Tnf* and *Tnfsf11* enriched in peripheral iNKT1b cells (**Fig. 3A**). These differences in steady-state expression of *Ifng* and *Il4* transcripts between iNKT1b and iNKT1c subsets prompted us to evaluate the production of these cytokines upon stimulation. *In vitro* stimulation with PMA and ionomycin induced strong IFN-γ and IL-4 responses by iNKT1b cells in the spleen and liver (**Fig. 3B, C**). iNKT1c cells had a more robust IFN-γ production than iNKT1b cells but produced little to no IL-4 (**Fig. 3B, C**). Following intravenous αGalCer injection *in vivo*, we found similar (spleen) or increased (liver) proportions of IFN-γ^+^ iNKT1b cells than IFN-γ^+^ iNKT1c cells, but only a few IL-4^+^ iNKT1c cells (**Fig. 3B, 3C**). Overall, these results indicate that iNKT1b and iNKT1c cells have distinct cytokine responses following stimulation, with iNKT1c cells producing lower amounts of IL-4 and IFN-γ upon *in vivo* antigen stimulation.

**Figure 3.**
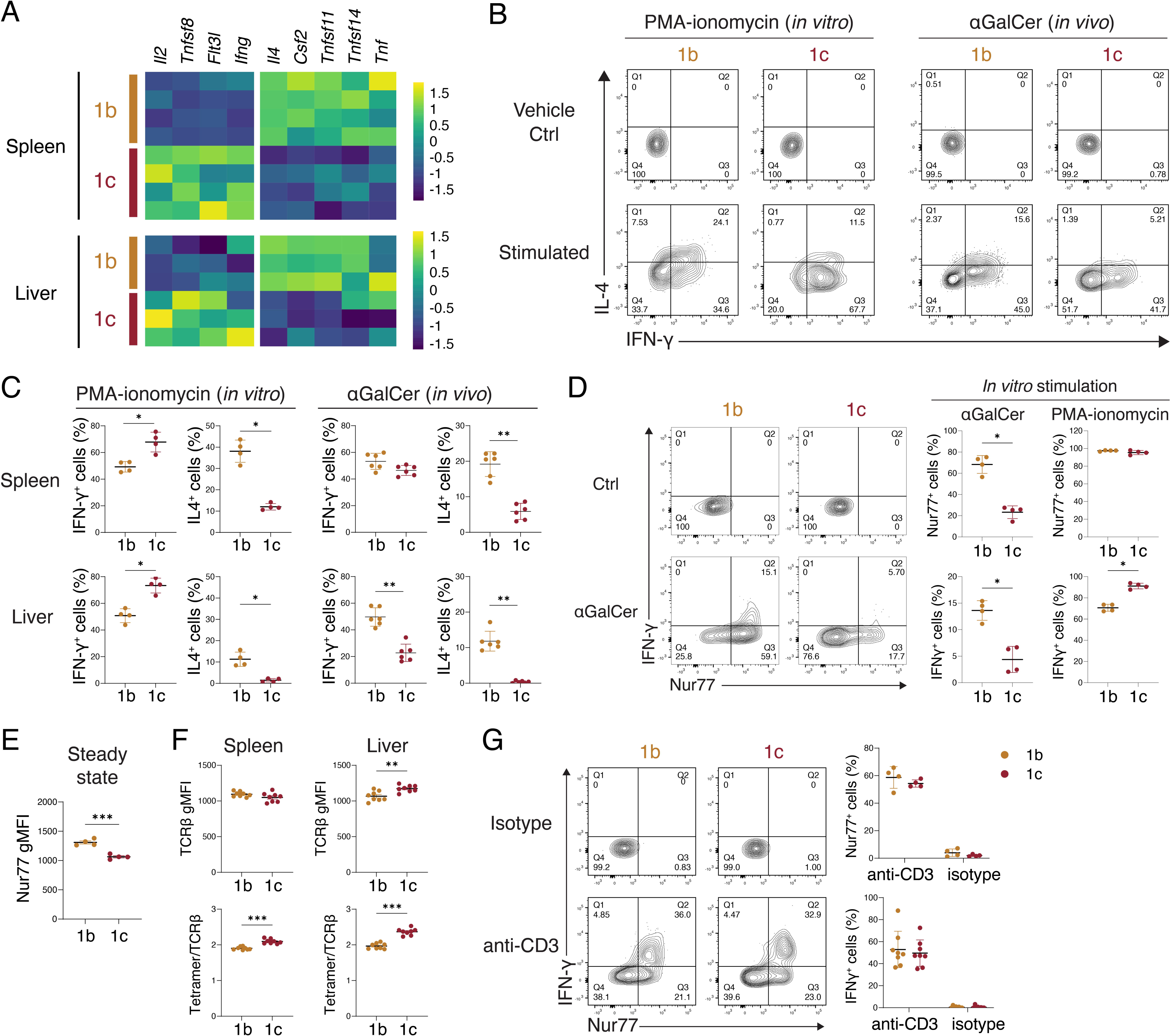
Weaker αGalCer-mediated IFN-γ response by iNKT1c cells. **A)** Heatmap showing cytokine DEGs for iNKT1b (1b) and iNKT1c (1c) cells sorted from the spleen and liver and analyzed by bulk RNA-seq. **B, C)** Splenocytes and liver leukocytes were stimulated *in vitro* with PMA and ionomycin or vehicle control for 4 h followed by intracellular cytokine staining for FACS analysis (left panels). B6 mice were injected intravenously with 0.5 μg αGalCer or vehicle control, then cytokine expression in iNKT cells from the spleen and liver was evaluated by FACS (right panels). Representative plots showing the expression of IL-4 and IFN-γ (B) and frequencies of IL-4^+^ and IFN-γ^+^ iNKT1b and iNKT1c cells are shown (C). **D)** Splenocytes from B6 mice were stimulated *in vitro* with 75 ng/mL αGalCer for 8 h or PMA and ionomycin for 4 h. Representative plots (left) and the frequency of iNKT1b and iNKT1c cells expressing Nur77 and IFN-γ (right) are shown. **E)** Geometric mean fluorescence intensity (gMFI) of Nur77 in iNKT1b and iNKT1c cells at steady state. **F)** gMFI and geometric mean of TCRβ and the tetramer/TCRβ staining ratio for iNKT1b and iNKT1c cells at steady state. **G)** Splenocytes from B6 mice were cultured in plates coated with 10 μg/mL anti-CD3 for 6 h. Representative plots of Nur77 and IFN-γ expression by iNKT1b and iNKT1c cells (left) and frequency of Nur77^+^ and IFN-γ^+^ iNKT1b and iNKT1c cells (right). Data shows the mean values ± SD. *** p<0.001, ** p<0.01, * p<0.05, Mann-Whitney test (C, D, E, F), Multiple t-test with Holm-Šídák correction (G).

The *in vitro* splenocyte stimulation with αGalCer described in the previous section resulted in a lower proportion of Nur77^+^ iNKT1c cells compared to iNKT1b cells (**Fig. 2D**), suggesting that iNKT1c cells may have lower sensitivity to antigen-mediated stimulation, which could explain their lower IFN-γ production *in vivo*. We confirmed that *in vitro* stimulation with αGalCer led to lower proportions of IFN-γ^+^ and Nur77^+^ iNKT1c cells compared to iNKT1b cells, while PMA-ionomycin stimulation led to higher frequency of IFN-γ^+^ iNKT1c cells and similar proportion of Nur77^+^ from both subsets (**Fig. 3D**). In addition, Nur77 expression at steady state was lower in iNKT1c cells than in iNKT1b cells (**Fig. 3E**). This lower response to antigen-mediated stimulation was not due to lower TCR expression or lower αGalCer-CD1d tetramer binding avidity by iNKT1c cells (**Fig. 3F**). Finally, both iNKT1b and iNKT1c cells responded similarly (in terms of IFN-γ^+^ cells and Nur77^+^ cells) to plate-bound anti-CD3 antibodies (**Fig. 3G**). Therefore, the lower iNKT1c response to αGalCer was only observed in the presence of antigen-presenting cells (APC) within splenocyte cultures, which suggested that the activation of iNKT1c cells by cognate antigens may be regulated by co-signals.

### Cytotoxic iNKT1c cell response is regulated by activating and inhibitory NK receptors

To delve into the co-signals that may regulate iNKT1c cell activation and response, we first investigated the role of signaling lymphocyte activation molecule family (Slamf) receptors, as iNKT1c cells show higher expression of 2B4 (Slamf4, encoded by the *Cd244a* gene) and both iNKT1b and iNKT1c cells express *Slamf7*. However, *in vitro* stimulation of splenocytes from *Cd244^-/-^* or *Slamf7^-/-^* mice with αGalCer did not result in an increased percentage of Nur77^+^ iNKT1c cells compared to littermate controls (**Sup. Fig. 2A, B**).

Next, we investigated the role of Natural Killer (NK) receptors, as several have been shown to regulate TCR-mediated responses ^37–39^. Many genes encoding for activating (e.g., *Klrb1c*, *Klrk1*, *Ncr1*, etc.) and inhibitory (e.g., *Klrc1*, *Klra9*, etc.) NK receptors were enriched in iNKT1c cells compared to iNKT1b cells (**Fig. 4A**). We next assessed whether TCR-mediated response in iNKT1c cells could be increased by blocking the inhibitory NK receptor NKG2A, which has been shown to regulate iNKT cell responses ^40,41^. The addition of neutralizing anti-NKG2A antibodies to αGalCer-stimulated splenocytes resulted in a modest but significant increase of IFN-γ-producing iNKT1c cells, but not IFN-γ-producing iNKT1b cells (**Fig. 4B**). Neutralizing NKG2A also increased Nur77 expression in both populations (**Fig. 4B**). Conversely, iNKT1c cells produced higher IFN-γ upon stimulation with activating anti-NK1.1 antibodies compared to iNKT1b cells (**Fig. 4C**). In addition, stimulation with IL-15 increased the expression of NK1.1 by both subsets (**Fig. 4D**), which could indicate another mechanism by which IL-15 increases the cytotoxic capacity of iNKT1c cells. Together, these results suggest that iNKT1c cell activation is regulated by activating and inhibitory NK receptor engagement.

**Figure 4.**
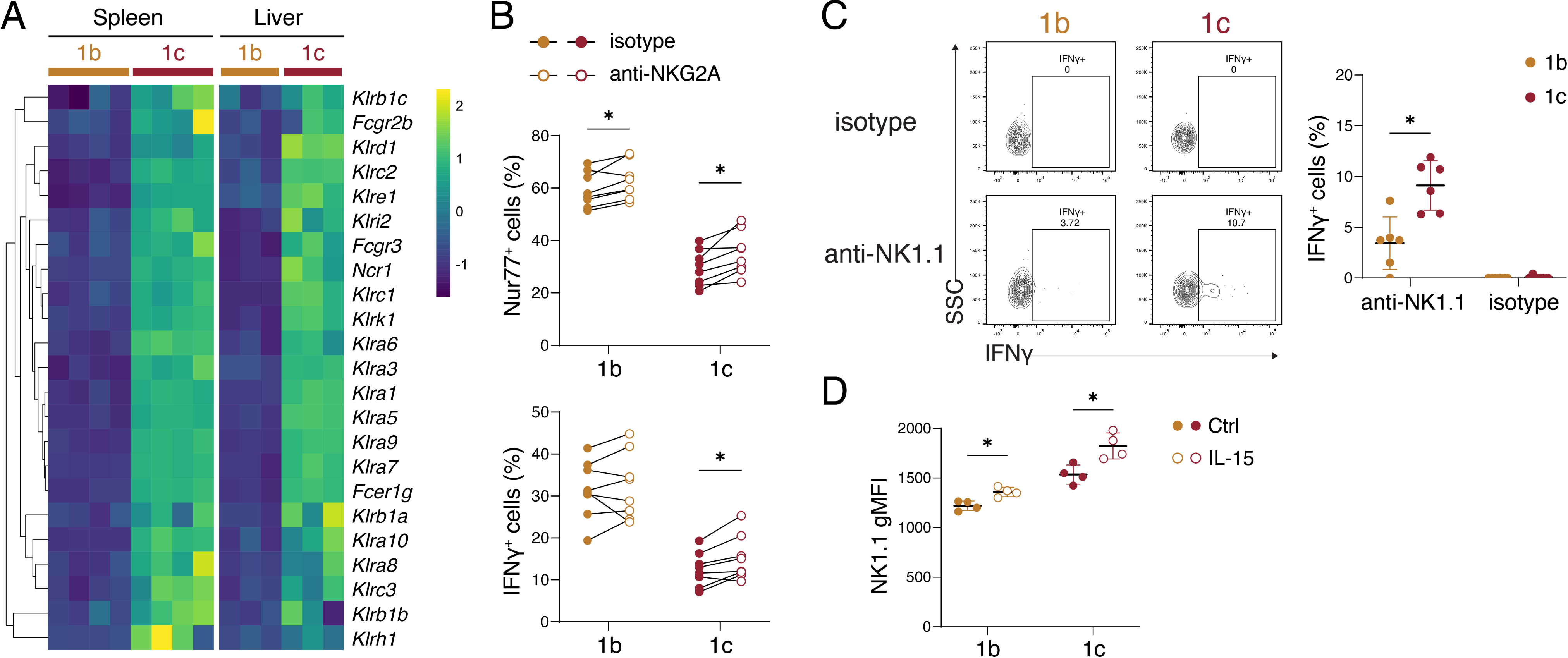
iNKT1c cell response is regulated by inhibitory and activating NK receptors. **A)** Heatmap showing the relative expression of genes encoding for NK receptors in iNKT1b (1b) and iNKT1c (1c) cells obtained by bulk RNA-seq. **B)** Splenocytes from B6 mice were stimulated with αGalCer in presence of 20 μg/mL blocking anti-NKG2A/C/E or the isotype control. Frequencies of Nur77^+^ and IFN-γ^+^ iNKT1b and iNKT1c cells are shown. **C)** Splenocytes from B6 mice were stimulated with plate-bound anti-NK1.1 (10 μg/mL) for 6 h. Frequency of IFN-γ^+^ iNKT1b and iNKT1c cells is shown. **D)** NK1.1 expression by iNKT1b and iNKT1c cells from spleen stimulated or not with 100 ng/mL IL-15 for 8 h. Data shows mean values ± SD. * p<0.05 Multiple t-test with Holm-Šídák correction.

### Antigen/CD1d-independent tumor killing activity by cytotoxic iNKT1c cells

Our data suggest that iNKT1c cells may be capable of recognizing tumor cells through NK receptors, perhaps even in the absence of TCR-mediated antigen recognition. To test this hypothesis, we assessed their ability to kill B16F10 melanoma cells expressing very low (B16F10-GFP) or high (B16F10F10-CD1d-GFP) levels of CD1d (**Fig. 5A**). FACS-purified iNKT1b and iNKT1c cells were expanded for a week in the presence of IL-2 (widely used for expansion of immune cells, including iNKT cells^42^) and IL-15 (shown to promote iNKT1 homeostasis^43,44^) and were then cultured with B16F10-GFP or αGalCer-loaded B16F10-CD1d-GFP cells using different effector-to-target (E:T) ratios. Of note, iNKT1c cells expanded more than iNKT1b cells (**Fig. 5B**). iNKT1c cells killed αGalCer-loaded B16F10-CD1d-GFP cells with greater efficiency than iNKT1b, at all E:T ratios (**Fig. 5C-E**). In addition, only iNKT1c cells were able to kill CD1d-negative/low B16F10-GFP cells (**Fig. 5C-E**). Anti-CD1d neutralizing antibodies did not impact the ability of iNKT1c cells to kill B16F10-GFP cells, confirming that their killing was not dependent on the low level of CD1d expression (**Fig. 5F**). Finally, we injected iNKT cell-deficient *Traj18^-/-^* mice intravenously with B16F10-CD1d-GFP cells to induce metastatic melanoma and adoptively transferred purified and *in vitro* expanded iNKT1b or iNKT1c cells, followed on the next day by a single intravenous αGalCer injection. Adoptive transfer of iNKT1c, but not iNKT1b, reduced the number of metastatic nodules in the lungs (**Fig. 5G**). We did not observe significant changes in the frequency or absolute numbers of conventional αβT cells, γδT cells, NK cells, B cells, or FoxP3^+^ CD4^+^ T cells in the lungs of treated mice compared to untreated controls (**Sup. Fig. 3A**). The frequencies and numbers of CD4^+^ and CD8^+^ cells and CD4/CD8 T cell ratio were also similar among groups (**Sup. Fig. 3B, C**). Moreover, no significant difference in PD1 expression by conventional T cells and γδT cells was observed (**Sup. Fig. 3D**). Of note, iNKT cells were detected in the lungs of all mice receiving iNKT1b or iNKT1c cells, but not in control mice (**Sup. Fig. 3E**). It is therefore likely that iNKT1c cells directly kill B16F10-CD1d-GFP cells, thereby limiting the number of lung nodules. Taken together, these results demonstrate that iNKT1c cells kill B16F10 tumor cells using CD1d-dependent and -independent signals, ultimately providing better tumor control.

**Figure 5.**
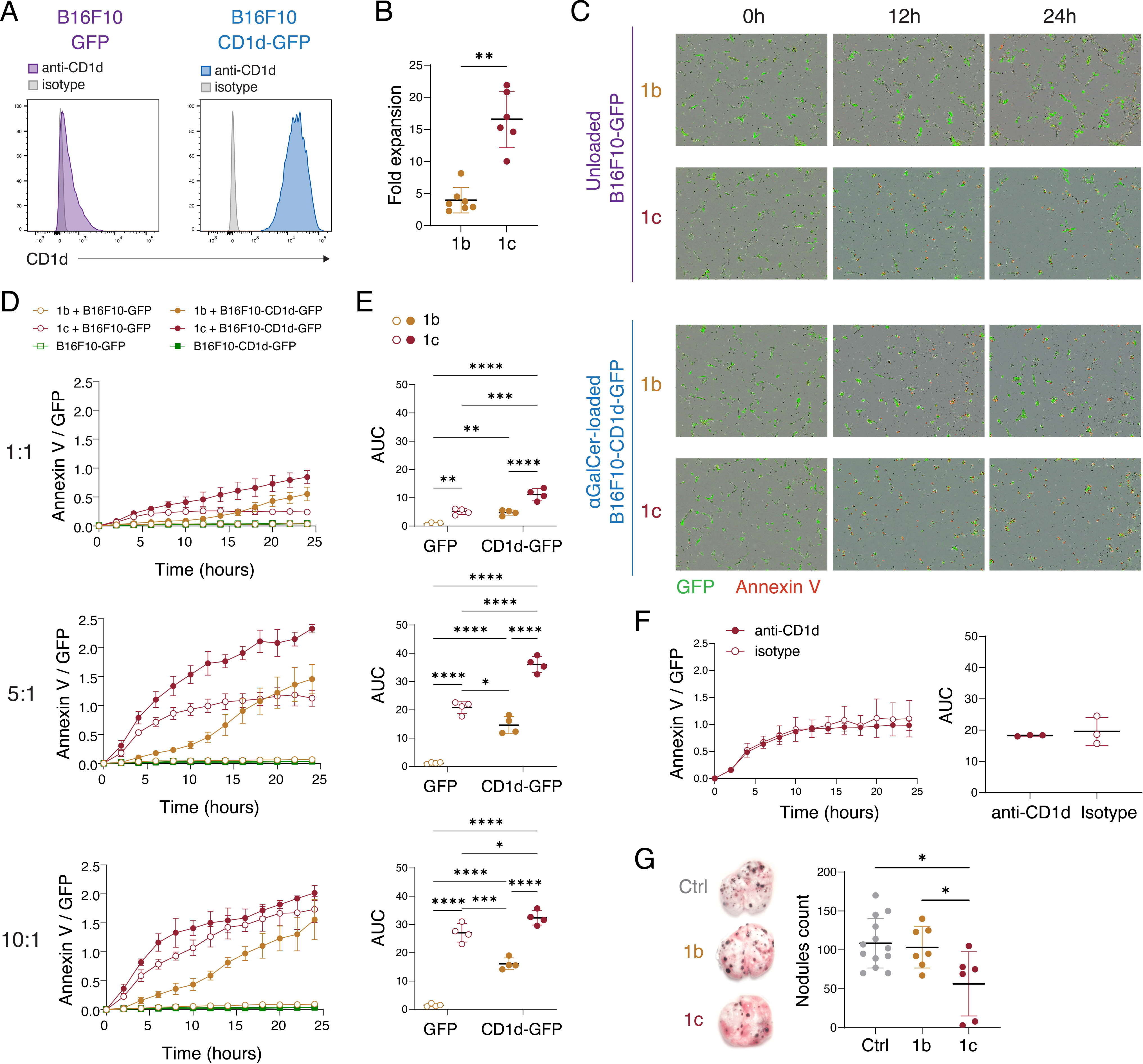
Antigen-dependent and -independent tumor cell killing by iNKT1c cells. **(A)** B16F10 melanoma cells were transduced to overexpress of CD1d and GFP (B16F10-CD1d-GFP), or only GFP (B16F10-GFP). Histograms show the expression of CD1d in both cell lines. **B)** iNKT1b (1b) and iNKT1c (1c) cells were purified from pooled livers of 6-to 8-week-old B6 female mice and expanded with plate-bound anti-CD3 in the presence of IL-2 and IL-15. Fold expansion of each population after 7 days is shown. **C)** Expanded iNKT1b and iNKT1c cells were co-cultured with αGalCer-loaded B16F10-CD1d-GFP or unloaded B16F10-GFP target cells in presence of Annexin V red reagent. Representative scan images at 10X obtained with the Incucyte are shown for co-cultures at 5:1 E:T ratio. **D)** Mean values ± SD of the Annexin V counts ratio over GFP^+^ counts at various E:T ratios. **E)** Area under the curve (AUC) calculated from the previous graphs for each co-culture. **F)** Unloaded B16F10-GFP target cells were pre-incubated with anti- CD1d (clone 1B1) or isotype control antibodies before co-culturing with iNKT1c cells at 5:1 E:T ratio. Annexin V/GFP ratio line graphs and the respective AUC plot are shown. **G)** 6- to 8-week-old female *Traj18^-/-^* mice were intravenously injected with 10^5^ B16F10-CD1d-GFP tumor cells. On the next day, 2 x 10^6^ *Traj18^-/-^* splenocytes alone (Ctrl) or in combination with 1.5 x 10^5^ expanded iNKT1b or iNKT1c cells were intravenously transferred. Mice were treated with 0.5 μg αGalCer one day after adoptive transfer. Representative images of lungs (left) harvested on day 15 after tumor cell injection and the metastatic nodules counts (right) are shown. Data shows mean values ± SD. **** p<0.001, *** p<0.001, ** p<0.01, * p<0.05, Mann-Whitney test (B, F), 2-way ANOVA with Tukey correction for multiple comparison (E), ANOVA with Tukey correction for multiple comparison (G).

### Retinoic acid signaling is required for the development of cytotoxic iNKT1c cells

Among the genes enriched in iNKT1c cells was *Rara* (**Fig. 6A**), which encodes for the retinoic acid receptor alpha (RARα), a nuclear transcription factor activated by retinoic acid (RA) that regulates the expression of RA target genes. Notably, RA is a vitamin A metabolite involved in many biological processes including T cell differentiation and NK cell function ^45–48^. To assess whether RA signaling regulated the development or homeostasis of iNKT1c cells, we analyzed mice with conditional expression of a dominant negative *Rara* (*dnRara*) gene targeted to the ROSA26 locus downstream of a *loxP-stop-loxP* (*lsl*) cassette, crossed with *Tbx21^RFP-Cre^* mice. In these mice, the RA-dependent activity of RARα is disrupted in T-bet-expressing cells, while its ligand-independent activity is retained ^46,49,50^. We found a drastic loss of iNKT1 cells and a concomitant increase in the frequency of iNKT17 cells in the thymus of *dnRara^lsl/-^ Tbx21^RFP-Cre/+^* mice (dnRara mice) compared to littermate control mice that were either *dnRara^lsl/-^ Tbx21^+/+^* or *dnRara^-/-^Tbx21^RFP-Cre/+^* (**Fig. 6B**). These phenotypes were more subtle in the spleen of dnRara-expressing mice and absent in the liver (**Fig. 6B**). Within iNKT1 cells, the frequency of iNKT1b cells was significantly reduced in all tissues analyzed, while the frequency of the closely related Tbet^+^ NK1.1^-^ iNKT cell population was significantly increased (**Fig. 6C**). Furthermore, iNKT1c cells, as well as GzmA^+^ iNKT1 cells were virtually undetectable in all tissues analyzed (**Fig. 6C, D**).

**Figure 6.**
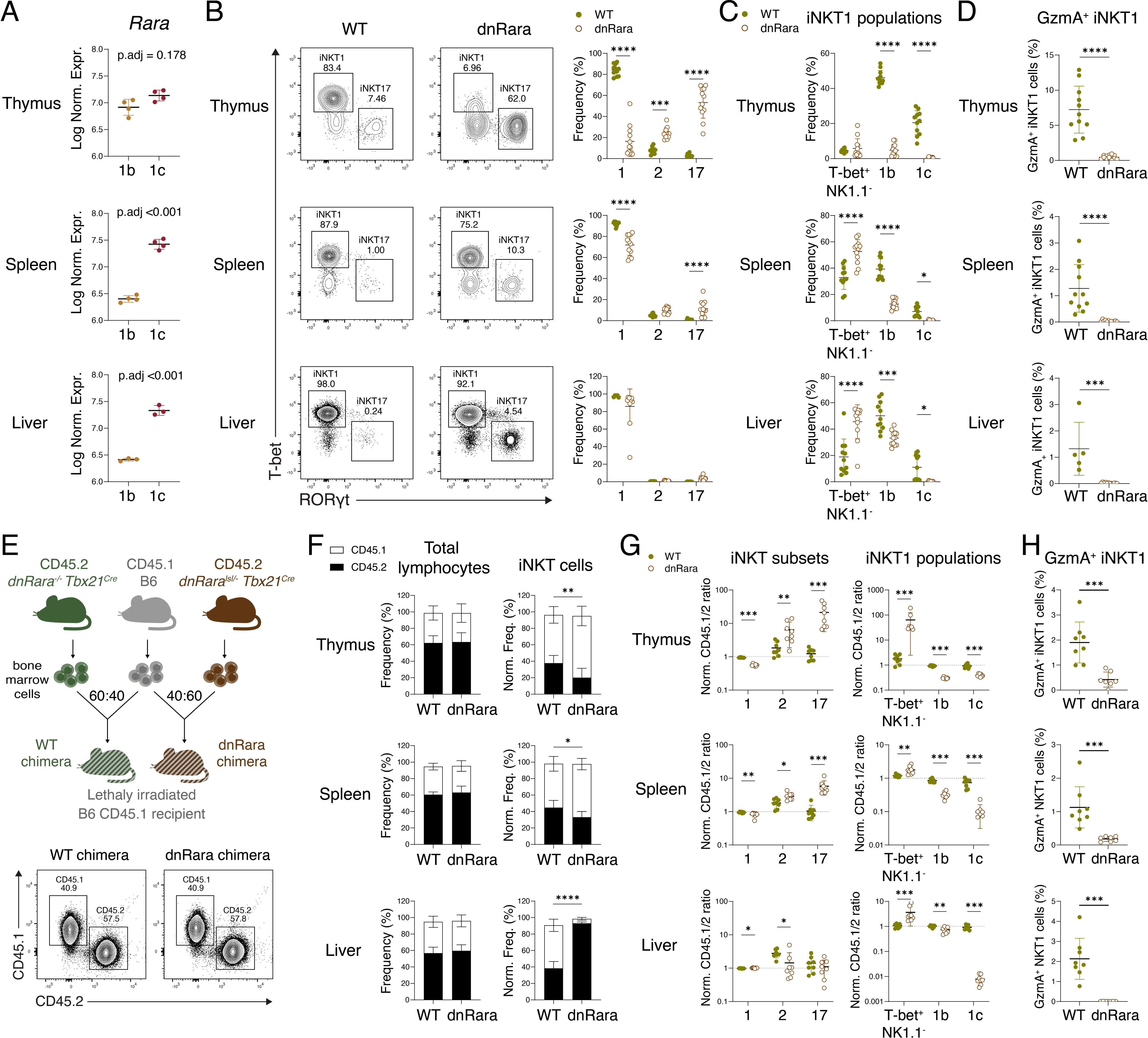
RA signaling is required for the generation of iNKT1c cells. **A)** iNKT1b (1b) and iNKT1c (1c) cells isolated from the thymus, spleen, and liver were analyzed by bulk RNA-seq. Log Normalized counts of *Rara* transcripts are shown in the plots. **B)** iNKT cells from 6-to 7-week-old female *dnRara^lsl/-^ Tbx21^RFP-Cre/+^* (dnRara) and *dnRara^lsl/-^ Tbx21^+/+^* or *dnRara^-/-^Tbx21^RFP-Cre/+^* littermate control mice (together referred to as WT) were analyzed by FACS. Representative FACS plots (left) and frequency (right) of iNKT subsets are shown. **C)** Frequency of T-bet^+^ NK1.1^-^, iNKT1b, and iNKT1c populations in the indicated tissue. **D)** Frequency of GzmA^+^ iNKT1 cells in the indicated tissues. **E)** Lethally irradiated sex-matched CD45.1 B6 mice were reconstituted with sex-matched *dnRara^lsl/-^ Tbx21^RFP-Cre/+^* and B6 CD45.1 (dnRara chimeras) or *dnRara^-/-^ Tbx21^RFP-Cre/+^* and B6 CD45.1 (WT chimeras) bone marrow cells (60:40 ratio). Representative FACS plots (bottom) show the chimerism for the spleen. **F)** Stacked plots show the mean proportion of CD45.1^+^ and CD45.2^+^ lymphocytes (left) and iNKT cells (right) in each chimera group. **G)** Normalized ratio of CD45.1^+^/CD45.2^+^ cells within each iNKT subset (left) and iNKT1 population (right) are shown. **H)** Frequency of GzmA^+^ iNKT1 cells in CD45.2^+^ cells from WT and dnRara chimeras. Data shows mean values ± SD error bars are shown. **** p<0.001, *** p<0.001, ** p<0.01, * p<0.05, Wald test with Benjamini and Hochberg correction for multiple testing (A), Mann-Whitney test (C, G), Multiple t-test with Holm-Šídák correction (B, F).

To determine whether this effect was due to cell-intrinsic signals, we generated mixed bone marrow chimeras by reconstituting lethally-irradiated and congenically-marked B6 CD45.1 mice with a 60:40 ratio of bone marrow cells from *dnRara^lsl/-^ Tbx21^RFP-Cre/+^* mice (CD45.2 allele) and B6 CD45.1 (dnRara chimeras), or a 60:40 ratio of bone marrow cells from *dnRara^-/-^ Tbx21^RFP-Cre/+^* (CD45.2 allele) and B6 CD45.1 (WT chimeras) (**Fig. 6E**). While the 60:40 CD45.2/1 chimerism was maintained in total lymphocytes found in the thymus, spleen and liver, a slightly lower proportion of dnRara iNKT cells compared to WT iNKT cells was observed in the thymus and spleen (**Fig. 6F**). However, there was a preferential accumulation of dnRara iNKT cells in the liver of these mice (**Fig. 6F**), which could indicate a role of RA signaling in iNKT cell homing or local proliferation. Moreover, we found a modest yet significant reduction in iNKT1 cell frequency in the thymus and spleen from dnRara chimeras (**Fig. 6G**, left panels). This suggests that both cell-intrinsic and -extrinsic factors are involved in the effect of RA signaling on iNKT1 cell differentiation, particularly in the thymus. Interestingly, similar frequencies of iNKT1 cells were found in the liver of WT and dnRara chimeras. Despite this, the frequency of iNKT1c cells were severely reduced in all tissues of dnRara chimeras, and especially in the liver (**Fig. 6G**, right panels). In addition, GzmA^+^ iNKT1 cells were greatly reduced in the thymus and spleen of dnRara chimeras, and almost absent in the liver (**Fig. 6H**). Together, these results indicate that RA has a complex and tissue dependent effect on the generation of iNKT1 cells. Specifically, cell-intrinsic RA signaling is essential for the generation or maintenance of cytotoxic GzmA^+^ iNKT1c cells.

### Cytotoxic human iNKT cells infiltrate non-small cell lung cancer primary tumors

We next assessed whether similar cytotoxic iNKT cell populations could be found in humans. Peripheral blood mononuclear cells (PBMCs) were analyzed for expression of molecules associated with the cytotoxicity (e.g., CD16, GzmB, GNLY), cell maturation (e.g., CD57), activation/exhaustion (e.g., PD-1), or molecules that have been previously proposed to correlate with a iNKT cell cytotoxic expression profile (e.g., CD94, 2B4) ^31,32,35^, using spectral flow cytometry. Unsupervised clustering analysis of the combined iNKT cells from healthy donors revealed 10 clusters of iNKT cells, some of which correspond to previously characterized populations (**Fig. 7A**). For instance, cluster 5 iNKT cells are characterized by the expression of CD62L and CD4, a phenotype characteristic of undifferentiated and highly proliferative iNKT cells (**Fig. 7B, C**) ^31,51–53^. Cluster 1 iNKT express CD4, low levels of CD161 and lack CD62L, and are likely another immature subset ^52–55^. Cluster 10 iNKT cells are CD62L^+^ CD4^-^ (**Fig. 7B, C**). Although such iNKT cells have previously been reported, their features remain poorly understood ^51,52,56^. Mature CD8^+^ CD161^+^ iNKT cells, which have been shown to appear later during ontogeny ^53,54,57^, constituted cluster 6 iNKT cells in our analysis. Furthermore, we found three clusters (2, 3, and 8) with a distinctive cytotoxic profile, characterized by expression of markers related to cytotoxic functions such as GzmB, granulysin (GNLY), and CD16. Among these, cluster 8 iNKT cells are characterized by the high expression of CD94, as well as 2B4, GzmB, and GNLY (**Fig. 7B**). These cells may correspond to a cluster of iNKT cells identified in previous scRNA-seq studies, which was enriched by the genes *KLRD1, GZMB,* and *GNLY* (encoding for CD94, GzmB, and GNLY, respectively). Cluster 3 iNKT cells contain CXCR6^+^ 2B4^+/low^ cells that express some levels of GzmB. CXCR6^+^ 2B4^+^ mouse iNKT cells have been previously described as highly cytotoxic in a recent study ^35^. Although the authors suggested that the same markers identify a cytotoxic population of human iNKT cells, we were not able to detect a unique cluster with those features. We found that clusters 3, 6, and 7 all co-expressed variable levels of CXCR6 and 2B4, but only cluster 3 showed GzmB expression (**Fig. 7B**). Finally, cluster 2 iNKT cells exhibited the highest expression of GzmB, GNLY, 2B4, and CD16, as well as variable expression of CD94 (**Fig. 7B**). These cells uniquely express high levels of CD57, and are CD4^-^ CD8^-^, or CD8^+^ (**Fig. 7B, C**). Although CD57^+^ iNKT cells have been reported before, CD57 expression was not associated with cytotoxicity ^58^. Consistent with previous findings, we found concomitant expression of RORγt and T-BET in cluster 3, 6, and 7 iNKT cells ^59^. However, cluster 2 CD57^+^ iNKT cells expressed the highest levels of T-BET and low or no RORγt (**Fig. 7B**). We did not find a distinct cluster expressing only RORγt. The proportion of each of the populations identified was highly variable among donors, and even undetectable in some donors. We considered the total number of cells in each cluster as well as their sample representation to maximize precision. Clusters 8 and 10 had less than 100 cells but included cells from more than 80% of the donors. On the other hand, cluster 9 consisted of only 26 cells from female donors and was only present in 25% of the donors (**Sup. Fig 4A, B**). Therefore, we excluded cluster 9 from further analysis. Furthermore, by analyzing individual samples we verified the preferential expression of GzmB and GNLY by CD57^+^ and CD94^+^ iNKT cells (**Sup. Fig. 4C, D**). As expected, only a minority of the CXCR6^+^ 2B4^+^ iNKT cells expressed those cytotoxic markers.

**Figure 7.**
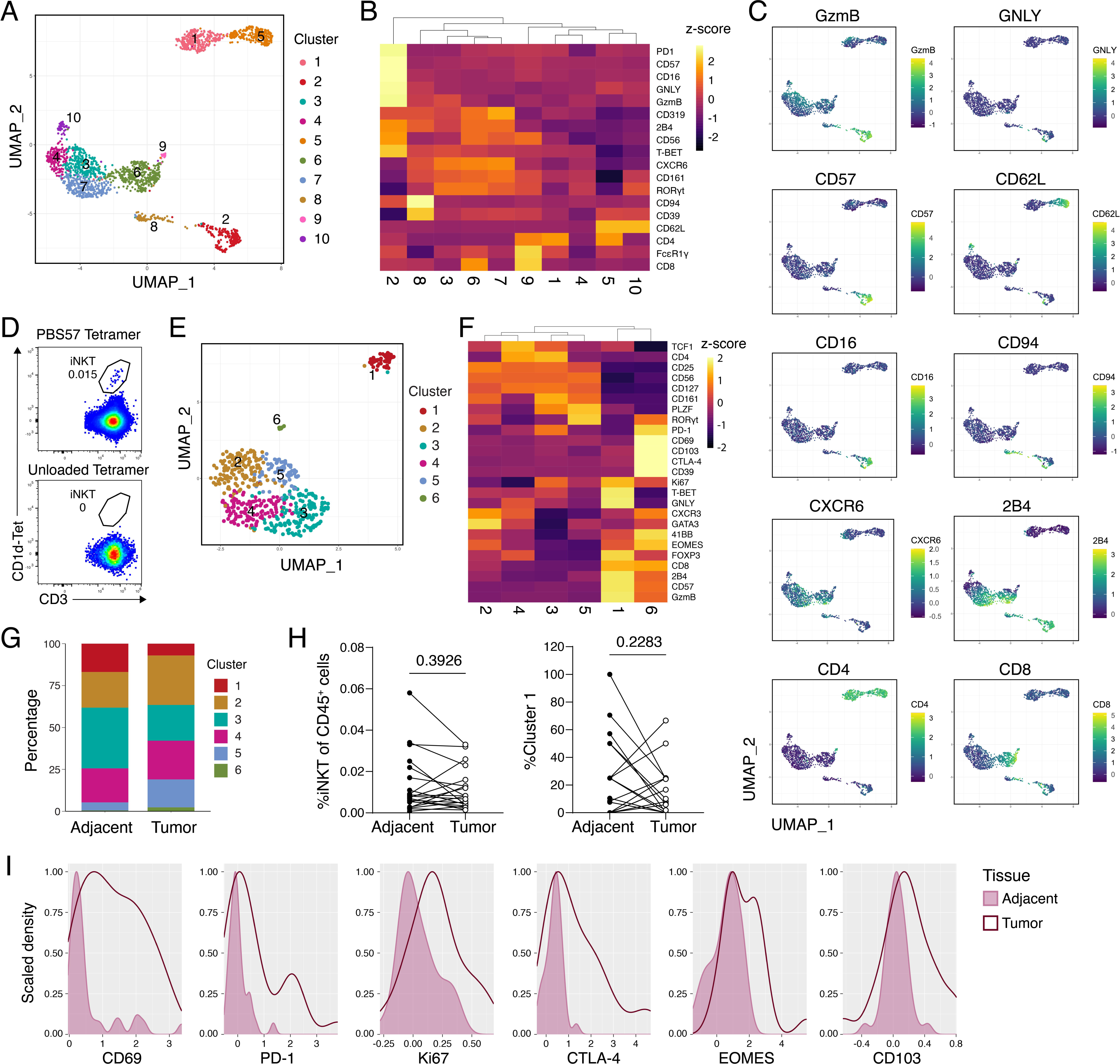
Human iNKT cells with cytotoxic profiles are found in peripheral blood and lung tumors. **A)** PBMCs from 14 healthy donors were isolated and analyzed by spectral FACS. Phenograph clustering analysis was performed on iNKT cells pooled from the 14 donors (200 iNKT cells per donor). UMAP dimensional reduction was performed to visualize the clusters. **B)** Heatmap of the scaled median expression of the markers in each cluster. **C)** Expression heatmap of selected markers that showed distinctive expression between clusters. **D)** Tumor and adjacent tissue were obtained from NSCLC patients and analyzed by spectral FACS. Samples were stained with PBS57-loaded or unloaded CD1d tetramers to control for iNKT cells gating. **E)** iNKT cells from matched tumor and adjacent tissues from 21 patients were concatenated and analyzed by phenograph for clustering. Identified clusters are shown in the UMAP plot. **F)** Heatmap of the scaled median expression of the indicated markers. **G)** Proportion of each cluster in tumor and adjacent tissue from the pooled data. **H)** Frequency of total iNKT cells (left) and cluster 1 iNKT cells (right) in each matched sample (Wilcoxon test). **I)** Density plots overlaying the expression of selected markers by cluster 1 iNKT cells in tumor and adjacent tissue (CV = 20.4).

Human iNKT cells can also be found in secondary lymphoid organs, but little is known about their phenotype in those tissues^60,61^. Moreover, the distribution of immune subsets in lymphoid tissues may not be mirrored in peripheral blood. Interestingly, CD57^+^ T-BET^hi^ GzmB^+^ GNLY^+^ iNKT cells were also detected in normal human spleen from a cadaveric donor (**Sup. Fig. 4E**). Thus, the presence of human iNKT cells with cytotoxic potential is not limited to peripheral blood.

As our mouse data demonstrated that iNKTc cells could infiltrate B16F10 melanoma tumors, we next assessed whether cytotoxic iNKT cells were present in human solid tumors. We obtained tumor and adjacent lung tissue samples from 21 non-small cell lung cancer (NSCLC) patients and analyzed iNKT cell phenotypes by spectral flow cytometry and unsupervised clustering analysis. Although iNKT cells were rare, we were able to identify a distinct cytotoxic population (cluster 1) of CD57^+^ T-BET^hi^ GzmB^+^ GNLY^+^ 2B4^+^ iNKT cells, reminiscent of cluster 2 iNKT cells found in PBMCs from healthy donors (**Fig. 7D-F**). Although this cluster contained less than 100 cells, these cells were found in more than 50% of the samples and were detected in all experimental batches (**Sup. Fig. 4F, G**). This population was found at similar frequencies within the tumor and in the adjacent tissue (**Fig. 7G, 7H**). These cells also appeared to express higher levels of Ki67, CD69, CTLA4, PD-1, CD103, and EOMES inside the tumor than in the adjacent tissue (**Fig. 7I**, **Sup.** Fig. 4H), possibly indicative of activation and exhaustion induced by the tumor microenvironment. In addition, CD57^+^ iNKT cells (cluster 1) expressed higher levels of FOXP3 compared to other iNKT cell clusters (**Fig. 7F**), which could result from exposure to the TGF-β-rich tumour microenvironment ^62^. These results indicate that CD57^+^ iNKT cells with high cytotoxic potential are found in peripheral tissues and infiltrate solid tumors.

### Cytotoxic human iNKT cells and precursors can be expanded *in vitro*

CD57 expression has been associated with NK and T cell terminal differentiation ^63–65^. Although CD57^+^ iNKT cells could exert potent anti-tumor responses, they may have limited persistence or renewal capacity. Therefore, we attempted to identify precursors to these cytotoxic CD57^+^ iNKT cells. First, we performed a Slingshot pseudotime analysis from the spectral flow cytometry data obtained from the PBMCs of healthy donors, setting the most immature CD62L^+^ CD4^+^ (cluster 5) iNKT cells as the root. The algorithm returned three branches that shared a common origin (cluster 5) as well as CD62L^-^ CD4^+^ iNKT cells (cluster 4) (**Fig. 8A, B**). One of the branches included all the cytotoxic iNKT cells (clusters 2, 3, and 8), and ended with CD57^+^ iNKT cells (cluster 2), indicating terminal differentiation. The second branch included clusters 3 and 7, and ended with cluster 6 CD8^+^ iNKT cells, which is consistent with the hypothesis that CD8 expression appears later in the ontogeny of iNKT cells ^53,54^ The third branch was short and ended with CD62L^+^ CD4^-^ iNKT cells (cluster 10). This was somewhat unexpected as CD62L is usually expressed by undifferentiated iNKT cells, and interestingly may suggest that CD62L can be re-expressed by more mature iNKT cells.

**Figure 8.**
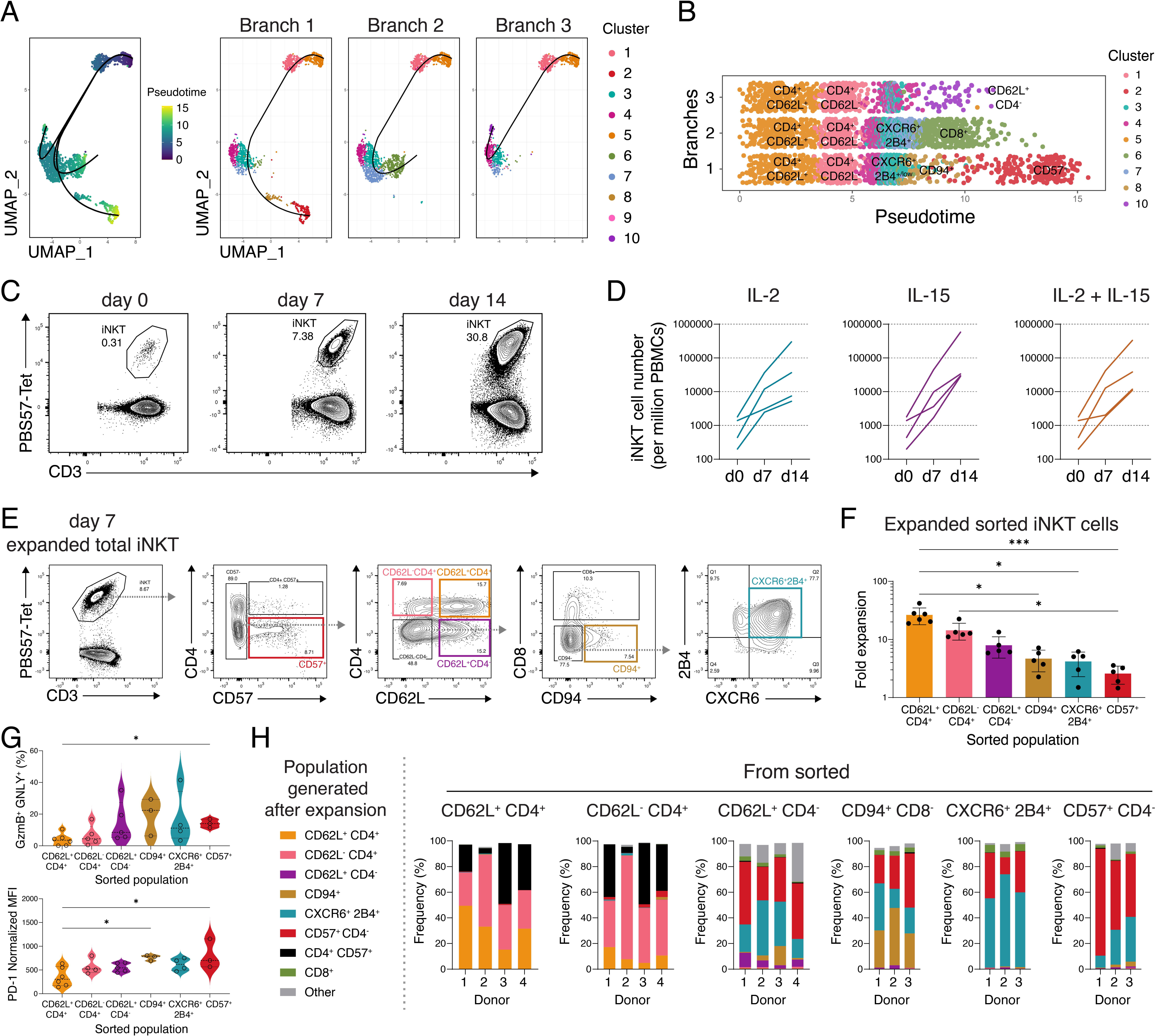
*In vitro* expansion and generation of human cytotoxic iNKT cells. **A)** iNKT cells from the PBMCs of healthy donors were analyzed by spectral FACS and phenograph clustering. Slingshot pseudotime analysis was performed placing cluster 5 (CD62L^+^ CD4^+^) iNKT cell at the origin. Three branches were obtained from the analysis. **A)** Linear representation of the pseudotime and the distinctive markers of the clusters are shown. **C)** Total PBMCs from healthy donors were co-cultured with αGalCer-loaded aAPCs at a 20:1 ratio in presence of 200 U/mL IL-2, 10 ng/mL IL-15, or a combination of 20 U/mL IL-2 and 10 ng/mL IL-15 (IL-2+IL-15) to expand iNKT cells for 14 days. Representative FACS plots of iNKT cells before and after expansion with IL-2+IL-15 are shown. **D)** Total number of iNKT cells obtained per initial million PBMCs at day 7 and 14. Each line represents a distinct donor. **E)** Six iNKT cell subsets were sorted after 7 days of culture. Representative plots show the gating strategy. **F)** The 6 purified subsets were expanded for 7 more days in the presence of IL-2+IL-15. The fold expansion is shown. **G)** Frequency of GzmB^+^ GNLY^+^ cells and PD-1 expression in each of the purified populations after expansion. **H)** Purified populations were analyzed by FACS after expansion using the same gating strategy shown in E to detect the populations generated *in vitro* post-expansion. Data mean ± SD is shown (F, G). *** p<0.001, * p<0.05, Kruskal-Wallis test and Dunn’s multiple comparison (F), Welch ANOVA and Dunnett T3 multiple comparison (G).

Next, we evaluated the capacity of selected iNKT cell populations to expand and differentiate in culture. To normalize activation between donors, PBMCs were stimulated with irradiated αGalCer-loaded K562 cells expressing CD1d, CD80, and CD83 as artificial antigen presenting cells (aAPC), as previously described ^66^, in the presence of IL-2 and/or IL-15. As IL-2 and IL-15, used either alone or in combination, induced similar proliferation of iNKT cells (**Fig. 8C, D**), and because IL-15 was found to promote iNKT cytotoxicity in mice, we used both cytokines for subsequent experiments. We expanded total iNKT cells from PBMCs for 7 days, FACS-purified six different iNKT cell populations based on our previous clustering analysis (**Fig. 8E**) and cultured them for 7 more days before analyzing them by FACS. As expected, the most undifferentiated CD62L^+^ CD4^+^ and the most mature CD57^+^ iNKT cells showed the highest and lowest expansion, respectively (**Fig. 8F**). At the end of the culture, CD57^+^ iNKT cells gave rise to the highest proportion of GzmB^+^ GNLY^+^ cells, which also expressed high levels of PD-1 (**Fig. 8G**). The six FACS-purified iNKT cell populations were able to generate other populations. For example, both CD4^+^ populations (either CD62L^+^ or CD62L^-^) gave rise to CD57^+^ CD4^+^ cells, which were barely detected before expansion and were not generated by any of the other populations (**Fig. 8H**), suggesting an alternative differentiation fate for CD4^+^ iNKT cells. CD62L^+^ CD4^-^ cells had intermediate expansion capacity and generated all the CD4^-^ populations including CD57^+^ iNKT cells. CD94^+^ iNKT cells generated CD57^+^ iNKT cells, as well as CD94^-^ CXCR6^+^ 2B4^+^ iNKT cells. CXCR6^+^ 2B4^+^ iNKT cells either remained unchanged or generated CD57^+^ iNKT cells. Finally, CD57^+^ iNKT cells mainly maintained their phenotype, but also gave rise to some CD57^-^ CXCR6^+^ 2B4^+^ iNKT cells (**Fig. 8H**). Overall, these results suggest that cytotoxic CD57^+^ iNKT are terminally differentiated and that they can be generated in culture from several other iNKT cell populations, including CD62L^+^ CD4^-^ iNKT cells, which show efficient expansion capacity and multipotency.

### Expanded human CD57^+^ iNKT cells exhibit direct cytotoxicity towards cancer cells

We expanded total iNKT cells from PBMCs for 7 days, FACS-purified the same six iNKT cell populations and cultured them for 7 more days, as described above, before testing their killing activity *in vitro* towards αGalCer-loaded B-cell lymphoblastoid C1R cells expressing nuclear GFP and transfected with CD1d (C1R-CD1d-NucGreen) or unloaded mock-transfected C1R (C1R-mock-NucGreen) cells. Real-time apoptotic ratios were obtained by measuring annexin V and GFP-expressing target cells with the Incucyte instrument. CD57^+^ iNKT cells showed significantly higher killing activity against αGalCer-loaded C1R-CD1d-NucGreen target cells compared to CD4^+^ populations (**Fig. 9A, B**). CD94^+^ cells showed a high cytotoxic activity, although the difference did not reach statistical significance. CXCR6^+^ 2B4^+^ iNKT cells displayed limited cytotoxicity. CD62L^+^CD4^-^ iNKT cells, which generate cytotoxic iNKT cells after expansion (**Fig. 8G**), killed αGalCer-loaded C1R-CD1d-NucGreen target cells with significantly higher efficiency than expanded CD62L^+^CD4^+^ iNKT cells (**Fig. 9A, B**). All iNKT cell populations similarly killed CD1d-negative C1R target cells at early time points (**Fig. 9C**). However, this was insufficient to prevent the outgrowth of these cells in these experimental conditions (**Fig. 9C**). Altogether, these findings indicate that CD57^+^ iNKT cells have the strongest capacity to kill tumor cells, with phenotypic and functional properties that overlap with iNKTc populations observed in mice.

**Figure 9.**
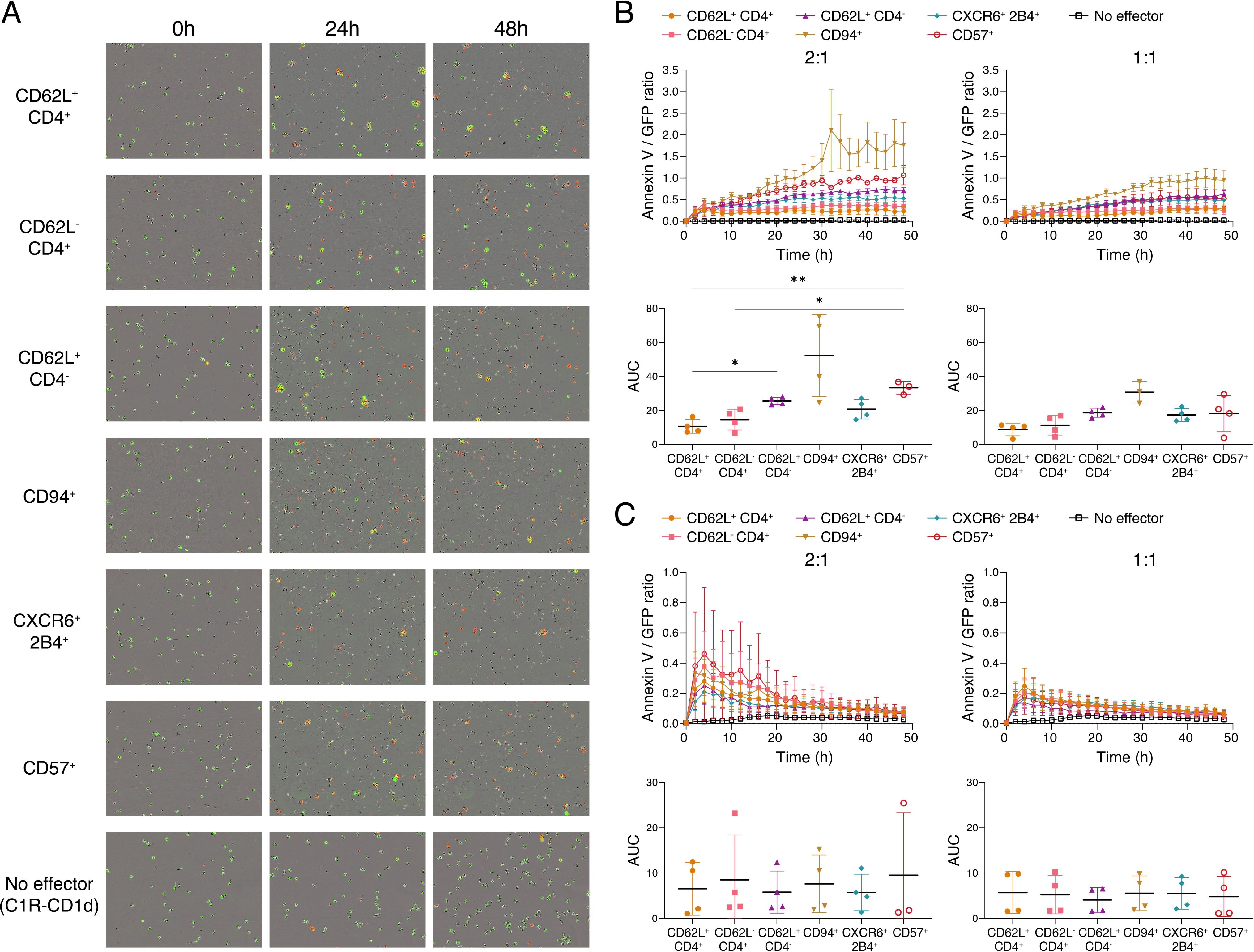
*In vitro* tumor killing activity of purified human iNKT cell populations. **A)** Six iNKT cell subsets were purified from 7 day-expanded iNKT cells, and further expanded for 7 days in presence of IL-2+IL-15 and co-cultured with αGalCer-loaded C1R-CD1d-NucGreen or unloaded C1R-NucGreen target cells in presence of Annexin V red reagent. Co-cultures were scanned every 2 h for a total of 48 h using the Incucyte instrument. **A)** Representative scan images (20X) of αGalCer-loaded C1R-CD1d-NucGreen co-cultures at 2:1 E:T ratio. **B)** Ratio of Annexin V/NucGreen counts (top panel) and area under the curve (AUC) (bottom panel) was calculated from αGalCer-loaded C1R-CD1d-NucGreen co-cultures at 2:1 (left) and 1:1 (right) E:T ratios. **C)** Ratio of Annexin V/GFP counts (top panel) and area under the curve (AUC) (bottom panel) was calculated from unloaded C1R-NucGreen co-cultures at 2:1 (left) and 1:1 (right) E:T ratios. Data mean ± SD is shown. ** p<0.01, * p<0.05, Welch ANOVA and Dunnett T3 multiple comparison.

## DISCUSSION

iNKT cells are recognized for their antitumor properties, and iNKT cell-based ACT to treat cancer and other diseases have recently entered clinical trials ^17–19,67^. However, whether cytotoxic effector functions against tumor cells are carried out by dedicated cytotoxic subsets remains elusive. Here, we identified and characterized iNKT cell subsets with *bona fide* cytotoxic effector functions and direct antitumor cytotoxicity in both mice and humans.

Using single cell transcriptomics, we and others recently showed that thymic iNKT1 cells are heterogenous and contain subsets with either helper or cytotoxic gene expression profiles, which we termed iNKT1b and iNKT1c, respectively ^29,30^. In the current study, we further characterized thymic and peripheral iNKT1c cells transcriptionally, phenotypically and functionally. These cells express T-bet and NK1.1, but unlike iNKT1b, they lack expression of Sca-1. iNKT1c cells have a cytotoxic effector program, produce granzymes A and B, and express a plethora of activating and inhibitory NK receptors. This subset may correlate with the 2B4^+^ CXCR6^+^ cytotoxic iNKT cell C2 subset recently reported by the Ikuta group ^35^. However, although these authors identified a human equivalent 2B4^+^ CXCR6^+^ iNKT cell subset, our unsupervised spectral flow cytometry analyses failed to identify such a unique subset in humans. In fact, 2B4 and CXCR6 are expressed at various levels by several iNKT cell subsets. 2B4^+^ CXCR6^+^ iNKT cells express low levels of granzyme B and granulysin, and display little to no cytotoxicity against B cell lymphoblastoid C1R tumor cells. By contrast, and in agreement with previous studies, we confirmed that CD94^+^ iNKT cells express high levels of cytotoxic molecules ^31,32^. In addition, we found that the most cytotoxic human iNKT cells uniquely express the sulfated oligosaccharide CD57, a marker expressed by highly cytotoxic and mature CD56^dim^CD16^+^ NK cells, memory-like NKG2C^+^ NK cells, and senescent CD8^+^ T cells ^63–65^. CD57^+^ iNKT cells express higher levels of T-BET, granzyme B, granulysin, CD16, CD94, 2B4, and CD56, but low levels of the inhibitory NK receptor CD161, compared to other iNKT cell subsets. Of note, CD57^+^ iNKT cells were previously identified in sarcoidosis patients ^58^, but CD57 expression was correlated with functional impairment (e.g.,, lower IFN-γ production), whereas the cytotoxicity of these cells was not assessed. Given that CD57 is a product of enzymatic activity, it was not detected in previous transcriptomic studies. Therefore, our study uncovered dedicated cytotoxic iNKT cell subsets in both mice and humans. We also demonstrated that their phenotypes differ between species.

Several studies have demonstrated that αGalCer mediated activation of iNKT cells controls the metastasis of B16F10 melanoma cells. This requires the transactivation of NK cells, which may or may not depend on IFN-γ production by iNKT cells ^5,68,69^. Moreover, metastasis prevention is best performed by CD4^-^ iNKT cells with low IL-4 production ^70^. Here, we showed that iNKT1c cells, but not iNKT1b cells, control lung metastasis of B16F10 cells *in vivo*. Although we have not assessed NK cell activation, this finding agrees with previously published data, as iNKT1c cells express less *Cd4* transcripts than iNKT1b cells and produce little to no IL-4 upon activation. We also demonstrated that mouse iNKT1c cells and human CD57^+^ iNKT cells directly kill B16F10 (and EL4) and C1R tumor cells *in vitro*, respectively. Besides direct cytotoxicity, it was shown that iNKT cells mediate antitumor immunity through the recruitment/activation of other innate and adaptive immune effector cells, the inhibition of immunosuppressive cells, and/or the promotion of antitumor immune memory ^67,71–74^. Whether iNKT1c cells can mediate these functions remains to be determined.

Whereas iNKT1b and iNKT1c cells were found in all lymphoid and non-lymphoid tissues tested, our transcriptomic analyses revealed that differences in gene expression between these two subsets were exacerbated in the spleen and liver, compared to the thymus. In agreement with this, it was recently shown that T-bet^+^ iNKT1 cells isolated from the thymus and the liver differed in their expression of genes associated with several biological processes, including cytokine signaling ^75^. In addition, the phenotype, TCR repertoire and antigen reactivity of iNKT cells is fine-tuned in peripheral tissues, both in mice and humans ^76^. These data suggest that tissue-specific cues (e.g.,, cytokines, antigens, microbial byproducts, hormones) may shape the phenotype and function of iNKT cell subsets. By extension, the tumor microenvironment could similarly impact the function of iNKT cells in cancer patients. We found mouse iNKT1c and human CD57^+^ cytotoxic iNKT cells within solid tumors. In mice, we found an increased proportion of cytotoxic iNKT1c cells in subcutaneous melanoma tumors, compared to other tissues in healthy mice. Whether this is the result of preferential recruitment, proliferation, survival and/or differentiation remains to be determined. Although mature iNKT cells are typically considered tissue resident, iNKT1c cells may be able to circulate between tissues, as indicated by their high expression of *klf2*, *S1pr1*, *S100a4*, *S100a6*, and low expression of tissue residency-related genes such as *P2rx7* (purinergic receptor P2X7). Similarly, the 2B4^+^ CXCR6^+^ C2 iNKT cell subset described by the Ikuta group was shown to circulate between tissues in parabiotic mice ^35^. In humans, cytotoxic CD57^+^ iNKT cells were found within primary NSCLC tumors and at the resection margin lung tissue at similar frequency.

In addition, intratumoral CD57^+^ iNKT cells appeared to be more activated/exhausted (higher expression of CD69, Ki67, PD-1, CTLA-4 and CD39), suggesting that they might be tumor reactive. Higher PD-1 expression within the tumors could make CD57^+^ iNKT cells susceptible to inhibitory signals from PD-L1-expressing tumor cells ^77,78^. This reinforces the rationale for combining anti-PD1 blockade with iNKT cell-based immunotherapies ^19,77,79,80^.

IL-15 plays crucial roles in the development and homeostasis of iNKT1 cells in mice ^43,44^. Here, we found that IL-15 increased iNKT1c cell granzyme B production and killing activity. In agreement with this, IL-15 production by thymic epithelial cells was required for the generation of cytotoxic 2B4^+^ CXCR6^+^ C2 iNKT cells ^35^. IL-15 also improved the *in vivo* persistence and antitumor activity of adoptively transferred human iNKT cells in tumor-bearing NSG mice ^81^. In addition, using mice expressing a dominant negative form of the RARα receptor, we demonstrated that retinoic acid (RA) signaling is necessary for the generation and/or maintenance of iNKT1c cells and GzmA-producing iNKT cells in general. Further investigation is required to decipher the role of RA signaling in the generation of mouse and human cytotoxic iNKT cells, as well as the mechanisms involved. Interestingly, a recent study demonstrated that TGF-β signaling promotes the differentiation of iNKT1 cells in the thymus, including 2B4^+^ iNKT1 cells, which may correspond to iNKT1c cells ^75^. However, the production of cytotoxic molecules (e.g.,, granzymes) and cytotoxicity were not evaluated in this study. A better understanding of the factors that control iNKT cell cytotoxic effector functions and their ability to kill tumor cells and more broadly promote antitumor immune responses will benefit the development of iNKT cell-based ACT to treat cancer.

Our data indicated that the response of mouse cytotoxic iNKT1c cells is regulated by TCR-independent co-signals, including activating and inhibitory NK receptors. For instance, we showed that CD94/NKG2A inhibits iNKT1c cell TCR signaling and their production of IFN-γ in response to αGalCer. In agreement with this, a previous study demonstrated that CD94/NKG2A interaction with its ligand Qa-1b inhibited cytokine production induced by αGalCer ^40^. In addition, antibody-mediated neutralization of NKG2A exacerbated αGalCer-induced liver injury^41^. Given that iNKT cells are autoreactive and that CD1d is ubiquitously expressed ^82^, it is perhaps not surprising that cytotoxic iNKT cells require additional checkpoints through inhibitory NK receptors, to prevent tissue damage. Other receptors, such as PD-1, may have a similar role ^83^, as we found that human cytotoxic CD57^+^ iNKT cells express higher levels of PD-1 than other subsets. On the other hand, we demonstrated that engagement of the activating NK1.1 (*Klrb1c*) receptor resulted in a modest but significant activation of iNKT1c cells. Mouse iNKT1c cells express a large panoply of genes encoding activating and inhibitory NK receptors, and our targeted analysis indicated that human CD57^+^ iNKT cells express relatively high levels of CD94, CD16, FcεR1γ, but low levels of CD161, compared to other subsets. It is therefore possible that several of these receptors regulate the activation of cytotoxic iNKT cells. For instance, human CD4^-^ iNKT cells express the activating receptor NKG2D and express perforin localized at the synapse with NKG2D ligand-expressing target cells ^39^. NKG2D engagement also potentiates iNKT cell activation ^39^. Characterizing the co-signals that regulate the function of cytotoxic iNKT cells will open further therapeutic possibilities. For example, the CRISPR-Cas9 gene editing of NKG2A in cytotoxic iNKT cells could unleash their cytotoxic effector functions in ACT, as it was recently demonstrated for NK cells ^84^. We found that mouse iNKT1c cells recognize and kill B16F10 tumour cells in CD1d-dependent and -independent manners. While the CD1d-independent killing of C1R cells by human iNKT cells was weak in our experimental conditions, several studies have shown that human iNKT cells can kill tumor cells in the absence of CD1d expression, through activating NK receptors such as NKG2D and/or CD226 (DNAM-1) ^85–88^. The identification of NK receptor and ligands involved in iNKT cell-mediated killing may ultimately enable the screening and stratification of tumors and patients for the rational deployment of iNKT cell-based ACT.

The success of ACT is intimately linked to the capacity of the transferred immune effector cells to expand and persist in the host ^89^. While CD57^+^ iNKT cells are highly cytotoxic, their limited proliferative capacity, at least *in vitro*, may limit their clinical use. Whether these cells can promote efficient and long-lasting antitumor responses will be the focus of future studies. However, we found that CD4^-^ CD62L^+^ iNKT cells exhibited a greater proliferative capacity and gave rise to cytotoxic iNKT cell subsets upon *in vitro* culture, which could kill C1R target cells. Moreover, our pseudotime analysis suggested that CD4^-^ CD62L^+^ iNKT cells developed from mature iNKT cells that re-expressed CD62L, suggesting that these cells are central memory-like. Work will be required to identify factors that preserve the persistence of adoptively transferred iNKT cells, particularly *in vivo*, while promoting their differentiation into cytotoxic effector cells. IL-21 may be an interesting candidate as it promotes the enrichment of all CD62L^+^ iNKT cells with a polarization towards a Th1 effector profile ^11,56^. In addition, a recent study demonstrated that human HSPC-derived CAR-iNKT cells engineered to express IL-15 persisted longer in NSG mice following adoptive transfer ^12^. In sum, we propose that CD4^-^ CD62L^+^ iNKT cells are an appealing candidate for ACT, which could be used alone or in combination with other short-lived but highly cytotoxic CD57^+^ iNKT cells.

This study expands our knowledge of the functional heterogeneity of iNKT cells in mice and humans. We described *bona fide* cytotoxic iNKT cell populations with the capacity to directly kill tumor targets, opening the door to harnessing cytotoxic iNKT cells in cancer immunotherapies. Moreover, we uncovered factors that can activate or inhibit the response of these cytotoxic populations, including NK receptors and cytokines. Ultimately, this work provides new insights to inform the rational design of iNKT cell-based cancer immunotherapies.

## METHODS

### Mice

C57BL/6J (B6) and B6 Cd45.1 mice were purchased from The Jackson Laboratory. *Traj18^-/-^* mice were kindly provided by Dr. Laurent Gapin (University of Colorado Denver, USA) ^90^. The *Tbx21^RFP-Cre^* mouse strain was generously shared by Dr. Alexander Rudensky (Memorial Sloan Kettering Cancer Center, USA) ^91^. The *Rosa26-dnRara^lsl^* mouse strain was generated and kindly donated by Dr. Cathy Mendelsohn (Columbia University, USA) ^49^. Cryopreserved sperm of *Klf2-Gfp* reporter mice were kindly provided by Dr. Stephen Jameson (University of Minnesota, USA) ^92^ and cryo-recovered at The Centre for Phenogenomics (TCP, Toronto). These mice were housed and bred under specific-pathogen free conditions (SPF) at the Division of Comparative Medicine animal facility (University of Toronto). All the procedures were approved by the Faculty of Medicine and Pharmacy Animal Care Committee at the University of Toronto (Animal use protocol #20012439). Sample tissues from *Cd244^-/-^* and *Slamf7^-/-^* mouse strains were a generous donation from Dr. Andre Veillette (University of Montréal, Québec).

### Human participants

All participants provided written informed consent before sample collection. Procedures were approved by the Health Sciences Research Ethics Board at University of Toronto (protocol # 00041425) and the Research Ethics Board at the University Health Network (UHN) (CAPCR protocols 15-9041, 21-5719). The study has been conducted in accordance with the principles stated in the Declaration of Helsinki.

### Reagents

The glycolipid antigen α-galactosylceramide (αGalCer, KRN7000) was purchased from Diagnocine (USA) as lyophilized powder. The powder was reconstituted by adding 0.05% Tween-20 in PBS, followed by heating at 80 °C and vigorous vortexing. Aliquots of 1 mg/mL αGalCer were stored at −20 °C. Biotinylated mouse CD1d monomers loaded with PBS57 (an αGalCer analog) ^93^ and unloaded monomers were generously provided by the National Institutes of Health (NIH) Tetramer Core Facility (Emory University, USA), and used to produce CD1d tetramers by combining the biotinylated monomers with fluorochrome-conjugated streptavidin at a molar ratio higher than 4:1 and in presence of 0.05% tyloxapol (Sigma-Aldrich).

### Cell lines

Mouse lymphoma EL4, EL4-GFP, and EL4-CD1d-GFP cell lines were maintained in Dulbecco’s Modiefied Eagle Medium (DMEM) supplemented with 10% Fetal Bovine Serum (FBS), 1 mM sodium pyruvate, MEM amino acids, MEM non-essential amino acids, 2 mM L-glutamine, 100 U/mL penicillin, 100 μg/mL streptomycin, and 50 μM β-mercaptoethanol. Mouse melanoma B16F10, B16F10-GFP, and B16F10-CD1d-GFP cells were cultured in DMEM supplemented with 10% FBS, 100 U/mL penicillin, and 100 μg/mL streptomycin. Human lymphoblast K562 cells expressing CD1d, CD80, and CD83 were kindly donated by Dr. Naoto Hirano (University of Toronto). Human B-cell lymphoblastoid C1R-CD1d (transfected for CD1d expression) and C1R-mock (mock-transfected) cell lines were generated and kindly provided by Dr. Steven Porcelli (Albert Einstein College of Medicine, USA). Human cell lines were maintained in complete RPMI (cRPMI), which consists of RPMI supplemented with 10% FBS, 1 mM sodium pyruvate, MEM amino acids, MEM non-essential amino acids, 2 mM L-glutamine, 100 U/mL penicillin, 100 μg/mL streptomycin, and 50 μM β-mercaptoethanol. C1R-CD1d-NucGreen and C1R-mock-NucGreen cells were generated by transducing C1R-CD1d and C1R-mock cells, respectively, and maintained in cRPMI containing 0.25 μg/mL puromycin.

### Mouse primary cell isolation

Mouse thymus, spleen, mesenteric lymph nodes (mLN), and inguinal lymph nodes (iLN) were harvested and homogenized with a plunger through a 40 μm cell strainer in Magnetic-Activated Cell Sorting (MACS) buffer consisting of 1X Phosphate Buffered Saline (PBS) containing 0.5% bovine serum albumin (BSA), and 2 mM EDTA into a 50 mL tube. After centrifugation at 1,500 rpm, 4 °C, the pellet from thymus, mLN, and iLN were resuspended in MACS buffer. Spleen cells were resuspended in 1 mL red blood cell (RBC) lysis buffer (Sigma-Aldrich), incubated at room temperature for 6 min and washed with MACS buffer. Then the leukocyte pellet was resuspended either in MACS buffer for staining or cRPMI for functional assays.

Mouse livers were perfused with cold PBS before harvesting and homogenized through cell strainers as described for the other tissues. Pellets from liver were resuspended in 10mL PBS, transferred to 15 mL tubes, and centrifuged once again. Then the pellet was resuspended with 8 mL of 33% Percoll (Sigma-Aldrich or Cytiva) in PBS and centrifuged at 1,300 x g for 20 min at room temperature, and with the brake off to isolate the leukocytes. The pellet was washed once with PBS and RBC lysis was performed as described for spleens. Cells were resuspended either in MACS buffer for staining or cRPMI for functional assays.

Mouse lungs were perfused with cold PBS through the heart before harvesting. Then, lungs were minced and transferred to 50 mL tubes containing 2 mL of digestion buffer (1 mg/mL type IV collagenase and 0.1 mg/mL DNase [Sigma-Aldrich] in Dulbecco’s PBS [DPBS] with calcium and magnesium). Tissues were digested for 45 min at 37 °C with constant shaking at 190 rpm. Digested lung pieces were homogenized through cell strainers while washing with MACS buffer. After centrifugation, lymphocytes were isolated by density gradient using 40% Percoll overlaid on top of 80% Percoll in PBS and spinning at 2,500 rpm, 20 min, room temperature, with the brake off. The interphase containing lymphocytes was collected and washed with PBS. The final pellet was resuspended in MACS for subsequent staining.

Subcutaneous B16F10 melanoma tumors were collected, minced, and digested as described for lungs samples. After filtering through cell strainers and washing with MACS, leukocytes were isolated by density gradient using 33% Percoll as previously described for liver samples. The cells were washed with PBS and resuspended in MACS for subsequent staining.

### Human peripheral blood mononuclear cell isolation

After informed consent, blood from healthy donors (7 female, 7 male, 20-40 years old) was collected through phlebotomy into Vacutainer™ tubes (BD) containing acid citrate dextrose (ACD). Peripheral blood mononuclear cells (PBMCs) were isolated using Sepmate tubes (Stemcell) following the instructions provided by the manufacturer. Briefly, blood was first diluted 1:1 in PBS containing 2% FBS (2%FBS-PBS), then 30-34 mL of diluted blood was layered on top of 15 mL Lymphoprep gradient medium (Stemcell) in a 50 mL Sepmate tube. Tubes were centrifuged for 10 min at 1,200 x g, room temperature, with the brake on. After centrifugation, the top layer was poured into a new 50 mL tube, topped up to 40 mL with 2%FBS-PBS, and centrifuged at 300 x g for 8 min at room temperature, with the brake on. Pellets from the same donor were pooled and washed once again with 2%FBS-PBS. Cells were then counted, resuspended in FBS containing 10% Dimethylsulfoxide (DMSO), and cryopreserved in liquid nitrogen until analysis.

### Human spleen preparation

A spleen was obtained from a healthy deceased donor (male, 72 years old). Pieces of approximately 1 cm^2^ were placed in 10 mL PBS containing 2% FBS and homogenized for 10min at room temperature using a gentleMACS dissociator (Miltenyi Biotec). Homogenized tissue was filtered through a 70 μm cell strainer and pooled. After RBC lysis, single cell suspensions were counted, resuspended in FBS containing 10% DMSO, and cryopreserved in liquid nitrogen until analysis.

### Human tumor sample preparation

Tumor and adjacent non-cancer tissue samples from 31 non-small cell lung cancer (NSCLC) patients (15 female, 16 male, 49-89 years old) were obtained from surgically resected lungs, as per UHN approved protocols. Tissue samples were then minced into fragments and enzymatically digested for 1 hour at 37°C into single-cell suspensions using the gentleMACS Octo dissociator and human tumor dissociation kit (Miltenyi Biotec) following the manufacturer’s instructions. Cells were resuspended with 12.5% human serum albumin (Gemini) in RPMI and 10% DMSO for cryopreservation in liquid nitrogen until analysis.

### Cell staining for flow cytometry and cell sorting

Single cell suspensions of mouse samples were stained for surface markers at 4 °C for 25 min in MACS buffer containing anti-CD16/CD32 diluted 1:200 to block Fc receptors, Live/Dead Aqua reagent (Invitrogen) diluted 1:1,000, and the desired fluorochrome-conjugated antibodies or tetramer at a dilution previously established by titration. For mouse tumor samples and samples prepared for sorting, the Fc block antibody was added 10 min before the rest of the antibody cocktail. For spectral FACS of human samples, the surface staining was split into 2 steps, each of them performed in MACS buffer with 10% Brilliant Stain Buffer Plus (BSB) (BD) and fluorochrome-conjugated antibodies at a dilution previously established by titration. Before staining, Fc block was performed by incubating cells for 10min with TruStain FcX (Biolegend) diluted 1:25 in MACS buffer. Then, cells were incubated with the first antibody cocktail for 20 min incubation at room temperature. After washing, staining with FVS780 (BD) or Live/Dead Aqua reagent diluted 1:900 in PBS was performed at room temperature for 15 min. Cells were then washed, stained with the second antibody cocktail for 25 min at 4 °C, and washed once again with MACS. After surface staining, cells were either resuspended in MACS for immediate analysis or continued the staining process for cytoplasmic markers and/or transcription factors. Fluorescence-minus-one (FMO) or fluorescent-minus-five (for spectral FACS) controls were included in the experiments.

For staining of cytoplasmic markers (cytokines), cells were fixed with Cytofix/Cytoperm buffer (BD Biosciences) for 30 min at room temperature. After washing with Cytoperm buffer (BD Biosciences), pellets were stained for 30 min at room temperature with the desired antibodies diluted in Cytoperm buffer. Cells were then washed with perm buffer and either resuspended in MACS buffer or continued the staining procedure for transcription factors.

For transcription factor staining, cells were fixed with FoxP3/transcription factor Fixation buffer (eBioscience) for 30 min at room temperature. Cells were then washed with Perm buffer (eBioscience) and stained for 30min at room temperature with the desired antibodies diluted in the same buffer. After washing with perm buffer, cells were resuspended in MACS buffer and either analyzed immediately or stored overnight at 4 °C before analysis.

### Flow cytometry and analysis

Samples were acquired for flow cytometry (FACS) analysis using the BD FACSymphony™ A5, BD FACSymphony™ A3, BD LSRFortessa™ X-20, BD LSRFortessa™, or Sony ID7000 analyzers. Performance QC and Daily QC were completed for the cytometers as recommended. For runs in BD analyzers, after optimizing the voltages for each panel, application settings were saved and verified using rainbow beads before each experiment to ensure consistency. Single stained Ultracomp beads (eBioscience) and SpectraComp beads (Slingshot) were used as compensation controls, except for the Live/Dead control, which consisted of cells with positive and negative populations in the same tube. Spectral unmixing values were calculated and autofluorescence subtracted by FACSDiva (BD) or ID7000 software (Sony), respectively. FCS files were analyzed using the FlowJo software. The coefficient of variation (CV) calculated for all the experiments was lower than 10%, unless otherwise specified in assays.

Cell sorting was performed using the BD FACSymphony™ S6 SE, BD FACSAria™ IIIu (operated by the facility staff), or BD FACSMelody™ (operated by user) sorters. CS&T and drop delay calibrations were performed before each sort. The sorting precision mode was set to “Purity”. Purified cells were collected either in 1.5 mL microtubes containing lysing buffer (for RAN-seq assays) or 5 mL polypropylene tubes containing 1.5 mL of medium with 30% FBS (for culture assays).

### Retroviral and lentiviral transduction

A mouse CD1d (mCD1d) construct was cloned into a pMSCV-IRES-GFP II (pMIG II) vector donated by Dr. Dario Vignali. Phoenix retrovirus producer cells were co-transfected with the packaging plasmid pCL-Eco (from Inder Verma) and the mCD1d-pMIG II construct or the pMIG II empty vector using the JetPrime transfection reagent (Polyplus) following the manufacturer’s instructions. At 24 h and 48 h of the transfection, the retrovirus-containing supernatants were collected, filtered, and stored at −80°C until use. The mCD1d-pMIG II and pMIG II retrovirus were used to transduce mouse EL4 and B16F10 cell lines. For an efficient transduction of B16F10 cells, the transient expression of the mouse cationic amino acid transporter 1 (mCAT1) was required, as mCAT1 is a cell surface receptor for ecotropic retrovirus. Thus, one day before the retroviral transduction, B16F10 cells were transfected with a mouse mCAT1construct using the JetPrime reagent. EL4 and mCAT1-expressing B16F10 cells were spinfected with the retrovirus by centrifuging them at 2,000 x g at 32 °C for 90 min in the presence of 8 μg/mL polybrene. Transduced cells defined as CD1d^+^ and/or GFP^+^ were isolated using a BD FACSMelody™ sorter. The stable expression of the transgenes was confirmed by FACS after several passages.

Incucyte NucLight Green Lentivirus Reagent (EF1a, Puro) (Sartorius) was used at a multiplicity of infection (MOI) of 6 to transduce C1R-CD1d and C1R-mock cells in the presence of 8 μg/mL polybrene and following the vendor’s instructions. After 24 h, medium was replaced with fresh cRPMI and the culture continued until confluence was reached. Then, transduced cells were selected by adding 1 μg/mL puromycin to the culture for 6 days. The expression of nuclear GFP in the transduced cells was verified by FACS and Incucyte SX5 (Sartorius). The resulting transduced cell lines were named C1R-CD1d-NucGreen and C1R-mock-NucGreen.

### RNA isolation and library preparation

All samples were obtained from 4 female 7-week-old *Tbx21^RFP-Cre^* (C57BL/6 background) mice. Thymic iNKT cells were magnetically enriched using the EasySep™ Mouse CD8α Positive Selection Kit II (Stemcell) with the modified protocol indicated by Stemcell to deplete CD8α^+^ cells (https://www.stemcell.com/using-easysep-positive-selection-kits-for-cell-depletion.html). Enriched cells from thymus, as well as splenocytes, and liver cells were stained and iNKT cell populations were sorted using the BD FACSAria™ IIIu sorter (for spleen) or BD FACSMelody™ sorter (for thymus and liver). 400 to 10,000 cells from each population were sorted into 1.5 mL tubes with 500 μL of RLT buffer (Qiagen). 5 μL of β-mercaptoethanol (BioUltra, Sigma-Aldrich) was added to each tube after sorting and samples were stored at −80 °C until the day of RNA extraction. RNA was extracted from thawed samples using the RNeasy Plus Micro Kit (Qiagen) according to manufacturer’s instructions. Immediately after RNA extraction, cDNA was obtained, tagged with 8-nucleotide unique molecular identifiers (UMIs), and amplified using the SMART-Seq mRNA LP (with UMIs) kit (Takara) according to manufacturer’s instructions and adjusting the number of cycles depending on the input number of cells. Amplified cDNA was purified using NucleoMag NGS Clean-up and Size Select (Takara) with a 1:0.8 sample:beads volume ratio as per the manufacturer’s instructions. Eluted cDNA was assessed for quality and quantified with the Agilent High Sensitivity DNA Kit and using a Bioananlyzer 2100 (Agilent) instrument. A dilution with a fixed concentration of 0.9 ng/μL cDNA was made for each sample and libraries were prepared from the diluted cDNA using the SAMRT-Seq kit and the Unique Dual Index Kit (Takara). Indexed libraries were pooled and purified with the NucleoMag NGS Clean-up and Size Select (Takara) as indicated by the kit. The pooled library was quantified by Qubit 4 fluorometer (Life Technologies) and the quality was assessed with a Bioanalyzer.

### RNA sequencing

A dilution of the indexed cDNA library at 5 ng/μL was prepared and sent for sequencing to Novogene Corporation Inc. (CA, USA). After passing quality control, the pooled library was sequenced in a full lane NovaSeq X Plus (367.5G), considering 2% PhiX and 150 paired end reads. More than 50 million reads were obtained per sample with Q30 higher than 85%. BCL files were obtained for analysis.

### Transcriptomic analysis

Base Calling was performed using BCL2FASTQ 2.20.0.422. The count matrix was obtained using the Cogent™ NGS Analysis Pipeline v2.0.1 (https://www.takarabio.com). Specifically, the “demux” command was used for demultiplexing. Then, mapping and read count quantification were performed by the “analyze” command from the pipeline, using mm10 as reference genome and “SMARTSeq_FLA_UMI” as experiment type. Total transcript counts were analyzed using the DESeq2 package from R ^94^. First, data was filtered to include only reads with more than ten counts and present in more than three samples. After estimating size factors and dispersions, differential expression analysis with adaptive shrinkage of the log fold change ^95^ was performed and regularized log transformation of the normalized counts was calculated. Euclidean sample distance plots, principal component analysis (PCA), and heatmap plots were obtained using the log normalized data. Gene symbols were mapped from the ENSEMBL transcripts using the AnnotationDbi package and the org.Mm.eg.db annotation data from R.

Gene ontology enrichment analysis of the differentially expressed genes (DEGs) (log2 flold change ≥ 1, p adj. < 0.05) was performed to determine overrepresented biological processes using the clusterProfiler package with a 0.05 q value cutoff ^96^. Gene set enrichment analysis of the ENSEMBL normalized counts using custom gene sets was performed with the GSEA v4.3.2 software^33,34^, considering the gene_set permutation type, 1000 permutations, collapse to gene symbols, omitted features with no symbol match, weighted enrichment statistics, and Signal2Noise metric for ranking genes.

Custom gene sets were obtained from our published single cell RNA sequencing (scRNA-seq) data. The data was reanalyzed using the Seurat package workflow ^97^. First, the reads were filtered to exclude transcripts in less than 3 cells, cells with more than 5% mitochondrial genes, cells with less than 1,500 reads or less than 900 transcripts, mitochondrial genes, ribosomal genes. Doublets were detected and excluded after analysis with DoubletFinder package ^98^. Briefly, doublets were predicted by the algorithm considering the optimal pk estimated by the mean-variance normalized bimodality coefficient, default pN value, the first 50 principal components, 5% expected doublet proportion, and adjustment for homotypic doublet proportion. After doublet filtering, reads were log normalized, and highly variable genes (HVG) were obtained considering a mean cutoff >0.0125 and <3, dispersion >0.5, and 2,000 features. Then, HVG were used to perform PCA on scaled data. The first 50 PC were considered for the clustering analysis using the Louvain algorithm with a resolution of 1.5, which was determined by analysis with the package clustree ^99^. Clusters were annotated based on our published article ^29^.

Next, we obtained DEGs of the iNKT1b cluster contrasted to the iNKT1c cluster. The genes were filtered to include only significant DEGs (p adjusted <0.01) with at least a two-fold change (log2 fold change ≥ 1). Finally, from these filtered DEGs, a list of the upregulated genes (iNKT1b_vs_iNKT1c_UP) and a list of downregulated genes (iNKT1b_vs_iNKT1c_DOWN) were obtained, which were then used as our custom gene sets for GSEA of the sorted populations.

### Clustering and pseudotime analysis of human iNKT cells

Exported FCS files of lymphocytes from each batch of tumor samples were first subjected to batch normalization by the CytoNorm package in R ^100^. Flowjo was then used to analyze the normalized data, gate on iNKT cells and export this population as a new FCS file. Only samples from patients with detectable iNKT cells in matched tumor and adjacent tissues were considered for subsequent analysis (21/31 total patients, 11 female, 10 male, 49-85 years old). Exported FCS files of iNKT cells from tumor samples and PBMCs were used for clustering analysis. First, the FlowCore package ^101^ was used to obtain the expression matrices and metadata. Expression data was then transformed by the hyperbolic arcsine (arcsinh) function. After merging the data, relevant channels (excluding dump or other lineage markers) were considered to perform unsupervised clustering analysis using the Rphenograph package ^102^ (k=40 and k=50 for iNKT cells from PBMCs and tumor samples, respectively). Dimension reduction by uniform manifold approximation and projection (UMAP) was performed to visualize the clusters. Expression data and cluster information were used to perform pseudotime analysis using the Slingshot package ^103^. All the channels used for the clustering analysis (without dimensional reduction) were considered as dimensions for the slingshot analysis. Only a starting (root) cluster was indicated in the analysis, which was selected based on expression of key markers previously assigned to highly immature iNKT cells according to several studies. Branches were obtained and curves embedded in the UMAP dimensions for visualization.

### Antigen-loaded bone marrow derived dendritic cells

Femur and tibia were obtained from female 6-to 7-week-old B6 mice. After removing muscle and connective tissue from the bones, epiphyses were cut leaving openings to the bone marrow. Next, the opened bones were placed inside a P1,000 pipette tip. The tips were then placed inside a 50 mL tube containing 100 μL of MACS buffer and centrifuged at 1,500 rpm, for 5 min at 4 °C. The tips with residual bones were discarded, while the remaining bone marrow cells at the bottom of the tube were subjected to RBC lysis and washed with PBS. To differentiate bone marrow cells into DCs, we used the procedure described by Gebremeskel *et al.* with minor modifications ^69^. Briefly, cells were counted and cultured in 6-well plates (1.5 x 10^6^ cells/well) containing complete RPMI (cRPMI, RPMI medium supplemented with 10% FBS, 1mM sodium pyruvate, MEM amino acids, MEM non-essential amino acids, 2 mM L-glutamine, 100 U/mL penicillin, 100 μg/mL streptomycin, 50 μM β-mercaptoethanol) and 40 ng/mL GMCSF. After three days, 2 mL of the culture supernatant was removed and replaced with fresh cRPMI with 40 ng/mL GMCSF. On day 6, non-adherent and semi-adherent cells were collected from each well by pipetting. Cells were then washed, counted, put back to culture, and loaded with the antigen. In this case, the culture medium consisted of cRPMI with 20 ng/mL GMCSF and 0.5 μg/mL αGalCer or the corresponding volume of vehicle control. Cells were incubated in this medium overnight and harvested by pipetting on day 7. The resulting loaded and unloaded bone marrow-derived DCs (BMDCs) were washed three times with PBS, counted, and adjusted to the desired concentration.

### *In vivo* iNKT cell stimulation assays

Short term stimulation of iNKT cells was performed by injecting female 6-to 7-week-old B6 mice with 0.5 μg of αGalCer through the tail vein. Mice injected with vehicle alone were used as control. After 90 min, spleen and liver were harvested, cells were isolated, and stained for the evaluation of cytokine production by flow cytometry. For long term stimulation, mice were intravenously injected with 6 x 10^5^ αGalCer-loaded or unloaded BMDCs ^69^. Then, spleen and liver were harvested at days 3, 7, and 10 to evaluate granzyme production by iNKT cells using FACS.

### *In vitro* stimulation assays

In all stimulation assays, splenocytes and liver cells obtained from 6-to 8-week-old B6 female mice were cultured in 24-well plates containing cRPMI and the respective stimuli. Cells were stimulated with 100 ng/mL phorbol 12-myristate 13-acetate (PMA, Sigma-Aldrich) and 1 μg/mL ionomycin (Sigma-Aldrich). The corresponding volume of DMSO was used in unstimulated control cultures. Cells were incubated for a total of 4h with the last 3.5 h in the presence of protein transport inhibitors (Invitrogen). Cells were then harvested, washed, and stained for flow cytometry analysis. Antigen stimulations were conducted using 75 ng/mL αGalCer or vehicle control for a total incubation period of 8 h. Protein transport inhibitors were added for the last 4 h. For cytokine stimulations, cells were incubated for 6-8h in presence of 100 ng/mL IL-2, IL-12, IL-15, or IL-18 (Biolegend). For plate-bound antibody stimulation assays, wells were first coated overnight at 4 °C with anti-CD3ε (clone 145-2C11, Invitrogen), anti-NK1.1 (clone PK136, Biolegend), or the respective isotype control antibody at 10 μg/mL in PBS. Wells were then washed 2 times with PBS and once with cRPMI. Cells were added and cultured for 6 h with addition of protein transport inhibitors for the last 4 h. In some stimulation experiments, key receptors were blocked to evaluate their role. First, using 24-well plates, cells were plated in 250 μL of medium containing 5 μg/mL of anti-CD16/CD32 (clone 93) to block Fc receptors and incubated for 15 min at culture conditions. Then, 250 μL of medium containing the blocking antibody was added to the cultures to a final concentration of 25 μg/mL anti-CD1d (clone 1B1, Invitrogen), 20 μg/mL anti-NKG2A/C/E (clone 20d5, Invitrogen), or the respective concentration of the isotype control, followed by an incubation of 30 min. Next, 250 μL of medium containing 4X concentrated stimuli was added. After 2 h, an additional 250 μL of blocking antibody or isotype control antibody was added to the cultures at the same final concentrations described before. The culture then continued the incubation period as established for each stimulation assay.

### *In vitro* expansion of mouse and human iNKT cells

Cells pooled from 5-10 livers isolated from 7-week-old female B6 mice were stained and iNKT cell populations sorted using a BD FACSMelody™ sorter. Sorted iNKT cells were cultured in cRPMI in 96-well plates that were previously coated for 12 h at 4 °C with 5 μg/mL of anti-CD3ε (clone 145-2C11, Invitrogen) in PBS, washed twice with PBS and once with cRPMI. The culture medium was supplemented with 10 ng/mL IL-2 and 25 ng/mL IL-15 (Biolegend). On the next day, cells were transferred to new uncoated wells and cultured in cRPMI with 20 ng/mL IL-2 and 50 ng/mL IL-15. Every two days, ¾ of the culture supernatant was replaced by fresh medium with cytokines. On day 7, expanded iNKT cells were collected, washed, counted, and adjusted to the desired concentration for subsequent assays.

Artificial antigen presenting cells (aAPCs) used to expand human iNKT cells consisted of K562 cells expressing human CD1d, CD80, and CD83, loaded with 0.1 μg/mL αGalCer for 2 h, and γ-irradiated with a dose of 200 Gy ^66^. PBMCs were co-cultured with aAPCs at a 20:1 cell ratio in the presence of 200 IU/mL IL-2, 10 ng/mL IL-15, or a combination of 20 IU/mL IL-2 and 10 ng/mL IL-15. Cytokines were replenished every 2-3 days and cells were restimulated with aAPCs every 7 days. To expand distinct iNKT cell populations, an initial cycle of 7-day expansion from total PBMCs was performed before sorting. Purified iNKT cell populations were then expanded for an additional 7 days with aAPCs in the presence of cytokines as mentioned before.

### End-point specific killing assay

EL4-CD1d-GFP cells were loaded with 0.5 μg/mL αGalCer in cRPMI for 5 h. EL4-GFP cells were incubated with the vehicle control in cRPMI for 5 h. After incubation, cells were collected and washed three times with PBS. αGalCer-loaded EL4-CD1d-GFP cells and unloaded EL4-GFP cells were labelled with CellTrace Violet (CTV, Invitrogen) diluted in PBS at 0.25 μM and 2.5 μM, respectively. Both CTV-labelled target cells were counted and mixed at 1:1 ratio before co-culture with effector cells. To prepare effector cells, iNKT cell populations were sorted from 10 pooled female B6 mouse livers, followed by a 6 h incubation with 100 ng/mL IL-15 (Biolegend) or vehicle control in cRPMI. Effector cells were then co-cultured with the target cell mix at a 5:1 ratio in 96-well plates. Cultures of target cell mix without effector cells were included as controls for specific killing quantification. After 10 h, cells were collected stained with LIVE/Dead Fixable Aqua reagent and analyzed by flow cytometry. The frequency of CTV^hi^ (unloaded EL4-GFP cells) and CTV^low^ (αGalCer-loaded EL4-CD1d-GFP cells) was determined from pre-gated Live GFP^+^ cells. The percentage of specific cell killing was calculated using the formula below:

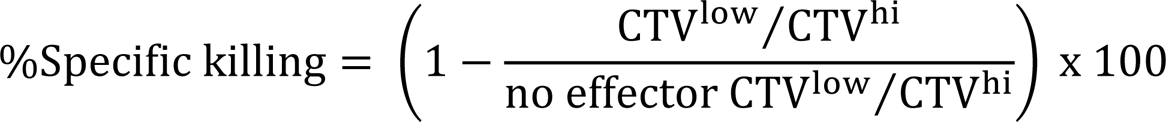

### Real-time killing assay

Adherent B16F10-CD1d-GFP and B16F10-GFP cells were seeded on 96-well plates at 2 x 10^3^ cells/well and cultured overnight. Then, the supernatant of B16F10-CD1d-GFP and B16F10-GFP cells was replaced by fresh medium containing 0.5 μg/mL αGalCer or vehicle control, respectively. Cells were incubated for 4h and washed once with warm cRPMI before the co-culture. For non-adherent cells, wells were pre-coated with 50 μL 0.01% poly-L-ornithine (Sigma-Aldrich) overnight. On the next day, the coating solution was discarded, and wells were washed first with PBS and then with cRPMI right before seeding the cells. To prepare the non-adherent target cells, C1R-CD1d-NucGreen and C1R-mock-NucGreen were incubated for 4 h with 0.2 μg/mL αGalCer or vehicle control, respectively. After washing twice with PBS and once with cRPMI, target cells were seeded on the coated plates (3,000 cells/well).

Expanded purified iNKT cell populations were used as effector cells, which were first harvested, counted, and diluted in cRPMI. Effector iNKT cells and target cells were co-cultured at different E:T ratios, in a total volume of 200 μL, and in the presence of Annexin V red reagent diluted 1:200 (B16F10 cell lines) or 1:400 (C1R cell lines). The plate was placed inside an Incucyte SX5 (Sartorius) and scans were taking after 30 min and every 2h for a total of 24-48 h. Two or four images per well were acquired at 10X or 20X augmentation, respectively.

For analysis of adherent cell killing, Top-Hat segmentation was used in the green (radius: 25 μm, threshold: 0.2, edge sensitivity: −80) and red (radius: 10 μm, threshold: 2.0, edge sensitivity: −35) channels, while the area was filtered to a minimum of 100 μm^2^ for both channels. For C1R cell lines, surface fit segmentation, as well as a minimum 10 μm^2^ area and 0.85 maximum eccentricity filters were applied for the green channel. Top-hat segmentation with 10 μm radius and 2 RCU threshold, and a minimum 100 μm^2^ area filter was applied for the red channel. Red/Green (Annexin V / GFP) count ratios were calculated for each well to assess apoptosis rate.

### B16F10 melanoma model and adoptive transfer

For the subcutaneous model, 6-to 8-week-old B6 female mice were subcutaneously injected on the left flank with 5 x 10^5^ B16F10 cells. Tumors were monitored daily with a caliper and the volume calculated using the formula: L x W^2^ x 0.52, where L is length and W is width. When tumors reached 1,000 mm^3^, mice were euthanized, tumors harvested and processed for flow cytometry analysis.

For the metastatic melanoma model, 6-to 8-week-old female *Traj18^-/-^* mice were intravenously injected with 10^5^ B16F10 cells in PBS. The next day, mice were intravenously injected with 1.5 x 10^5^ iNKT cell populations that were sorted from female B6 mice, expanded *in vitro*, and mixed with 2 x 10^6^ splenocytes from *Traj18^-/-^* female mice. As control, mice were injected only with 2 x 10^6^ splenocytes from *Traj18^-/-^* female mice. The following day, mice were intravenously injected with 0.5 μg of αGalCer. At day 15 post tumor cell injection, lungs were harvested, and the tumor nodules were counted under a stereoscope. Lungs were then processed for FACS analysis.

### Bone marrow chimeras

Cd45.1 B6 mice of 6 weeks old were exposed to γ irradiation for two cycles of 550 cGy. The mice were then reconstituted intravenously with 5 x 10^6^ bone marrow cells from sex-matched *dnRara^lsl/-^ Tbx21^RFP-Cre/+^* (dnRara mice) and Cd45.1 B6 mice at a 60:40 mix ratio, or from *dnRara^-l-^ Tbx21^RFP-Cre/+^* (WT mice) and Cd45.1 B6 mice at a 60:40 mix ratio. Reconstituted mice were treated with 2 g/L neomycin sulfate in drinking water for 14 days and returned to normal water for the rest of the experiment. Mice were euthanized 16 weeks after the reconstitution, and organs were harvested for FACS analysis.

### Statistical analysis

The precision in FACS experiments was assessed by the coefficient of variation (CV) considering a Poisson distribution. Thus, CV=100/√N, where N is the number of events in the rarest population. Statistical analyses were performed using R packages or the GraphPad Prism software. Mann-Whitney test was used for two-group univariate comparisons. Wilcoxon matched-pairs signed rank test was used for comparison of population in matched tumor and adjacent tissues. For experiments with more than two comparison groups or two explanatory variables, normality was first assessed by the D’Agostino-Pearson test and/or QQ plots, while homoscedasticity was assessed by Brown-Forsythe test or residuals plots. Based on the assumption, an appropriate statistical test was applied for each data set, which is specified in the legend of each figure. Parametric analysis included ANOVA test followed by post hoc analysis with Tukey correction, multiple t-test with Holm-Šídák correction, and Welch ANOVA tests followed by post hoc analysis with Dunnett T3 correction. Differential gene expression analysis was performed by the Wald test with Benjamini-Hochberg correction, implemented in the DESeq2 package. Non-parametric analyses included Kruskal-Wallis with Dunn’s multiple comparison test. Statistical significance was considered as p < 0.05.

## AUTHOR CONTRIBUTION

C.d.A.H. designed the research, performed and analyzed experiments and wrote the manuscript. S.W.Y.W., M.K. Z.L., J.A.M. and A.N. helped with some experiments. G.I. assisted on the analysis of the RNA-seq data. A.V., J.L.G., S.Q.C. M.S.T. and A.G.S. provided critical reagents or patient samples. T.B. and C.P. assisted on the design of the project and reviewed the manuscript. T.M. designed and supervised the research and wrote the manuscript.

## ACKNOWLEDGMENTS

We would like to thank the patients and their families for making this work possible. We would also like to thank the nurses, physicians and surgeons for efforts to obtain tissue samples. We are grateful to the healthy donors who participated in this study. This work was supported by the Canada Foundation for Innovation Physical Infrastructure Grant (John R. Evans Leaders Fund from the Canadian Foundation for Innovation application 29186), the Canadian Institutes of Health Research (application 363664) and a Tier 2 Canada Research Chair (file number 950-231786) to T.M. C.d.A.H. was supported by an Ontario Graduate Scholarship. The authors would like to thank Teresa Ciudad Garcia and Tracy McGaha for their assistance in experimental design and access to the Bioanalyzer.

## DATA AVAILABILITY

The RNA-seq data have been deposited on the Gene Expression Omnibus repository under the accession GSE272405.

