## Supplemental Figures for "*Bona fide* cytotoxic iNKT cells with superior antitumour responses identified in mice and humans"

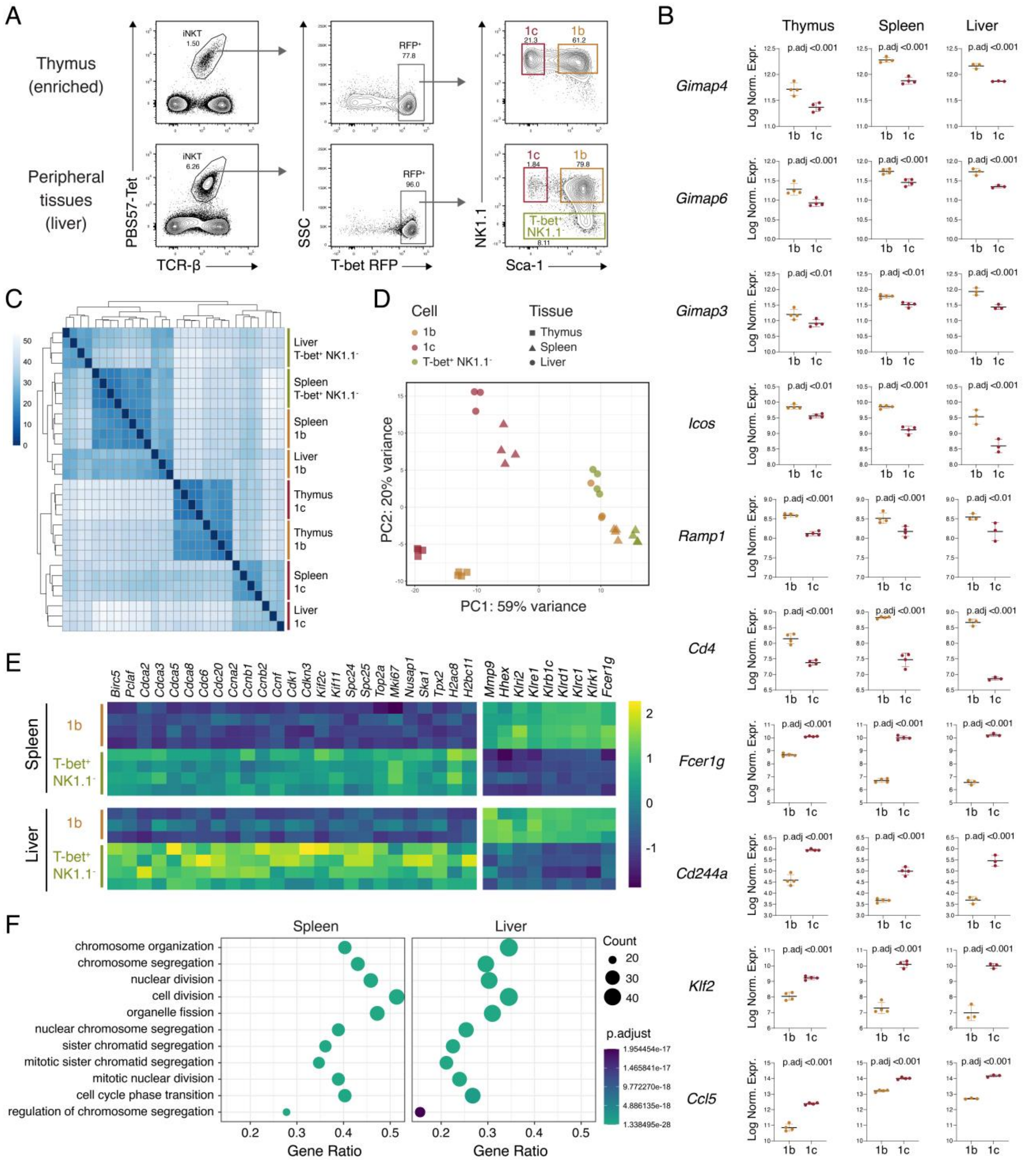


**Supplementary Figure 1. Transcriptomic features of iNKT1b, iNKT1c, and T-bet⁺ NK1.1⁻ iNKT1 subsets. A)** iNKT1b (1b), iNKT1c (1c), and T-bet⁺ NK1.1⁻ iNKT1 cells were sorted from the spleen and liver, and from magnetically enriched thymic iNKT cells of *Tbx21*$^{RFPCre}$ reporter mice for bulk RNA-seq analysis. Representative plots are shown. **B)**

Comparative plots showing the Log-normalized counts of some key genes with significant differential expression between iNKT1b and iNKT1c. Data mean ± SD is shown. Wald test with Benjamini and Hochberg correction for multiple testing. **C)** Sample distance plot obtained from the transcriptomic analysis of sorted iNKT1 populations. **D)** PCA plot of the transcriptomic data obtained from all the sorted populations. **E)** Heatmap of selected DEGs by peripheral iNKT1b and T-bet$^+$ NK1.1$^-$ iNKT1 cells. **F)** Biological processes overrepresented in T-bet$^+$ NK1.1$^-$ iNKT1 cells compared to iNKT1b cells from the spleen (left) and liver (right).

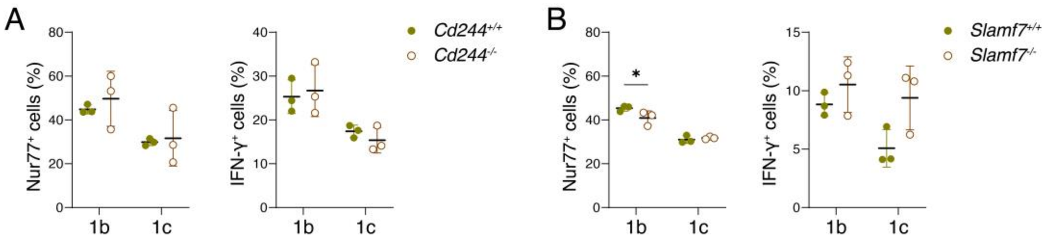


**Supplementary Figure 2. TCR-mediated IFN-γ response by iNKT1b and iNKT1c in Slamf4 and Slamf7-deficient mice. A, B)** Splenocytes from *Cd244*$^{-/-}$ (A) or *Slamf7*$^{-/-}$ (B) mice and their respective littermate controls were stimulated with 75 ng/mL αGalCer for 8 h and analyzed by FCM. Frequency of Nur77$^{+}$ and IFN-γ$^{+}$ cells in iNKT1b (1b) and iNKT1c (1c) populations. * $p<0.05$ Multiple t-test with Holm-Šídák correction.

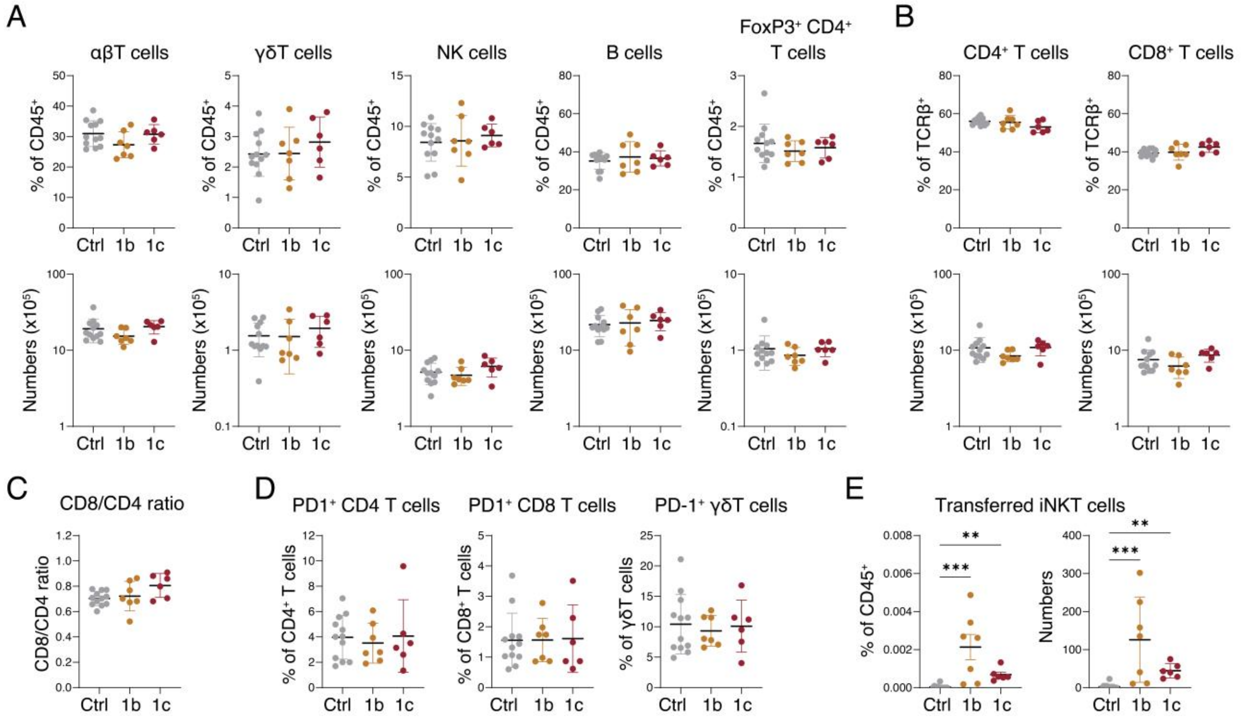

**Supplementary Figure 3. Lymphocyte profiling of lungs from metastatic melanoma-bearing mice following adoptive transfer of iNKT1b or iNKT1c cells.** 6- to 8-week-old female *Traj18*$^{-/-}$ mice were intravenously injected with $10^5$ B16F10-CD1d-GFP tumor cells. On the next day, 2 x $10^6$ *Traj18*$^{-/-}$ splenocytes alone (Ctrl) or in combination with 1.5 x $10^5$ expanded iNKT1b (1b) or iNKT1c (1c) cells were intravenously transferred. Mice were treated with 0.5μg αGalCer one day after adoptive transfer. **A)** Frequency (top) and absolute numbers (bottom) of the indicated lymphoid populations. **B)** Frequency and absolute numbers of CD4$^+$ and CD8$^+$ populations from conventional T cells. **C)** CD8$^+$/CD4$^+$ T cell ratios. **D)** Frequency of PD-1-expressing CD4$^+$ T, CD8$^+$ T, and γδ T cells. **E)** Frequency (left) and absolute numbers (right) of iNKT cells recovered from the lungs. Data shows mean values ± SD. *** $p<0.001$, ** $p<0.01$, * $p<0.05$, ANOVA with Tukey correction for multiple comparison or Kruskal-Wallis with Dunn's multiple comparison test.

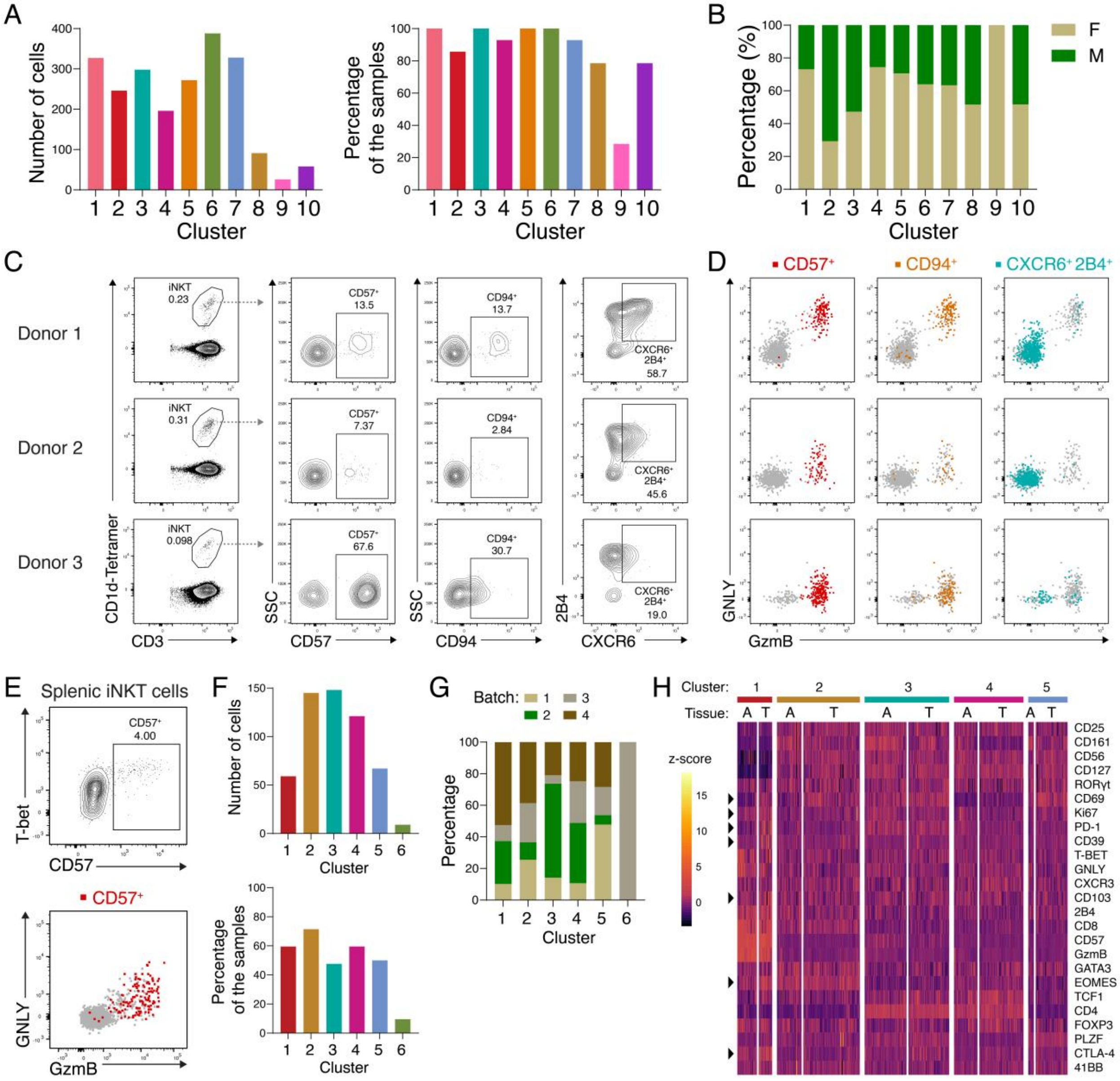


**Supplementary Figure 4. Analysis of human iNKT cell populations from PBMCs, spleen and lung tumors.** iNKT cells from PBMCs of healthy donors were analyzed by spectral FACS. **A)** After phenograph clustering analysis of 14 samples, the total number of events for each cluster and their representation across samples were calculated. **B)** Proportion of male and female donors for each cluster. **C)** Representative plot for the gating of CD57$^+$, CD94$^+$, and CXCR6$^+$ 2B4$^+$ iNKT cells (> 300 events) for three donors are shown. **D)** Expression of GzmB and GNLY for each of the gated populations. **E)** iNKT cells from the spleen of a healthy donor were analyzed by FACS. Expression of GzmB and GNLY by splenic $CD57^{+}$ T-$bet^{hi}$ iNKT cells is shown. **F)** Tumor and adjacent tissue were obtained from NSCLC patients and cells analyzed by spectral FACS. iNKT cells from matched tumor and adjacent tissue from 21 patients were concatenated and analyzed by phenograph for clustering. For each cluster, the total number of events and their representation across samples were calculated. **G)** For each cluster, the frequency of cells found in each experimental batch is shown. **H)** Heatmap of the scaled expression of markers in cells from the tumor and adjacent tissue for each cluster. Arrow heads point at some of the markers differentially expressed between tumor and adjacent tissue in cluster 1.
